# RNF43 is a gatekeeper for colitis-associated cancer

**DOI:** 10.1101/2024.01.30.577936

**Authors:** Alisa Dietl, Anna Ralser, Karin Taxauer, Theresa Dregelies, William Sterlacci, Mara Stadler, Roberto Olayo-Alarcon, Shushan Margaryan, Martin Skerhut, Tanja Groll, Katja Steiger, Dharmesh Singh, Xue Li, Rupert Oellinger, Roland Rad, Klaus Peter Janssen, Christian L. Mueller, Michael Vieth, Markus Gerhard, Raquel Mejías-Luque

## Abstract

Somatic mutations in the tumor suppressor Ring finger protein 43 (*RNF43*) were frequently found in colitis-associated cancer (CAC) and related to the duration of chronic inflammation, but their significance in inflammation and inflammation-associated carcinogenesis remained elusive.

We assessed the onset of *RNF43* mutations at different stages of human CAC development by exome sequencing, and comprehensively characterized *RNF43* loss-of-function-driven malignant transformation in mice by RNA sequencing, flow cytometry, immunohistochemistry, computational transcriptome-microbiome associations, and determined the underlying mechanisms by performing functional stem-cell derived organoid studies and fecal microbiota transfers.

Mutations in *RNF43* were frequent (12.9 %) in precancerous lesions of ulcerative colitis (UC) patients and eventually detectable in 24.4 % of CAC patients. In a bacterial-induced colitis mouse model, *Rnf43* mutations caused invasive colorectal carcinomas by aggravating and perpetuating inflammation due to impaired epithelial barrier integrity and pathogen control. We could demonstrate that *Rnf43* loss-of-function-mutations were even sufficient to cause spontaneous intestinal inflammation, resulting in UC-typical pathological features and subsequent invasive carcinoma development. In detail, mutant *Rnf43* impaired intestinal epithelial and particularly goblet cell homeostasis in a cell-intrinsic manner, and caused dysbiosis. The altered microbiota composition induced epithelial DNA damage and spontaneous mucosal inflammation characterized by TGF-ß-activating dendritic cells and pro-inflammatory (IL-17^+^, IL-22^+^, TNFα^+^) T cells. Over time, the continuous epithelial and goblet cell dysfunction, combined with pro-tumorigenic and pro-inflammatory microbiota, resulted in accumulated epithelial damage with transformation into inflammation-associated cancer in the presence of constitutive WNT signaling activation.

We identified mutant *RNF43* as susceptibility gene for UC and bona fide driver of CAC.

## INTRODUCTION

Patients receiving anti-inflammatory treatment are less likely to develop and die of CRC (1), stressing the general contribution of inflammation to CRC development. Accordingly, patients with inflammatory bowel disease (IBD) in particular are at high risk to develop colitis-associated cancer (CAC), and this risk directly increases with duration and extent of inflammation (2).

Evidence accumulates that IBD results from inadequate immune responses that are directed against microbiota and environmental factors in genetically susceptible individuals with epithelial barrier defects (3). Over 240 risk loci for IBD have been identified, many of which encode for proteins involved in microbiota-mediated immune signaling (4). The important contribution of the gut microbiota to inflammation and inflammation-driven tumorigenesis has been demonstrated in several mechanistic mice experiments which show that germ-free housing attenuates colitis in mice with depleted mucus barrier (5), while the presence of microbiota aggravates tumor burden in *Apc*^min^ mice (6). Alterations in the intestinal mucus barrier increase epithelial exposure to microbiota, metabolites, and food-born mutagens, resulting in spontaneous intestinal inflammation and epithelial damage (7). Mucus also serves as nutrient source for mucus-adherent microbiota and thus contributes to the selection of certain microbial species (8) that modulate mucus permeability (9), and sustain resistance against pathogen colonization (10). In line with this symbiont mucus-microbiota relationship, impaired goblet cell function with altered mucin biosynthesis is observed in IBD patients (11), and causes epithelial barrier impairment and dysbiosis with inflammation-driven cancer development in mice (12).

Somatic mutations in the WNT inhibitor Ring finger protein 43 (*RNF43*) were found in 11 % of CACs and related to chronic inflammation (13). In addition, *RNF43* expression was found to be strongly downregulated in CAC patients (14), and inactivating *Rnf43* mutations promoted tumor growth in a colitis-induced CRC mouse model (15). Yet, little is known whether and how mutated *RNF43* contributes to intestinal inflammation and inflammation-associated carcinogenesis.

In this study, we analyzed the frequency of *RNF43* mutations during the carcinogenic cascade in a human IBD cohort, and determined the effects of two *RNF43*-inactivating mutations on intestinal homeostasis in mice under basal and inflammatory conditions. Our results demonstrate that *RNF43* mutations arise early in IBD-lesions, and predispose to spontaneous inflammation-driven cancer by impairing epithelial and goblet cell homeostasis and shaping a DNA-damaging and Th17/Th22-inducing microbiota composition.

## RESULTS

### Mutations in *RNF43* are early events in the development of colitis-associated cancer

To investigate the potential significance of RNF43 as CAC driver mutation, we determined the prevalence of mutations in the *RNF43* exome at different stages of colitis-associated carcinogenesis in 102 colorectal tissue samples from 85 ulcerative colitis (UC) patients. *RNF43* was not mutated in the inflamed non-dysplastic mucosa, but in 7.14 % of low-grade dysplasia (LGD), 12.9 % of high-grade dysplasia (HGD), and 24.4 % of CAC samples (Figure 1A). We next related the *RNF43* mutation status to cancer stage, microsatellite status, cancer location and patient sex (Figure 1B). The prevalence of mutant *RNF43* increased with CAC progression, from 20 % in stage I CACs (2/10) to 37.5 % in stage II CACs (6/16) (Figure 1B). 72.7 % of *RNF43* mutant CACs (8/11) were microsatellite stable. Of note, these CACs were mostly located on the left side and showed primarily mutations in the *RNF43* RING domain (Figure 1B). The early and stepwise increasing prevalence of *RNF43* mutations in CAC precursors suggest that mutations in *RNF43* may drive the progression of non-cancerous lesions to CAC. This was supported by the observation that CAC patients with mutated *RNF43* tended to have a shorter transition time from IBD onset to CAC diagnosis (Figure 1C).

**Figure 1.**
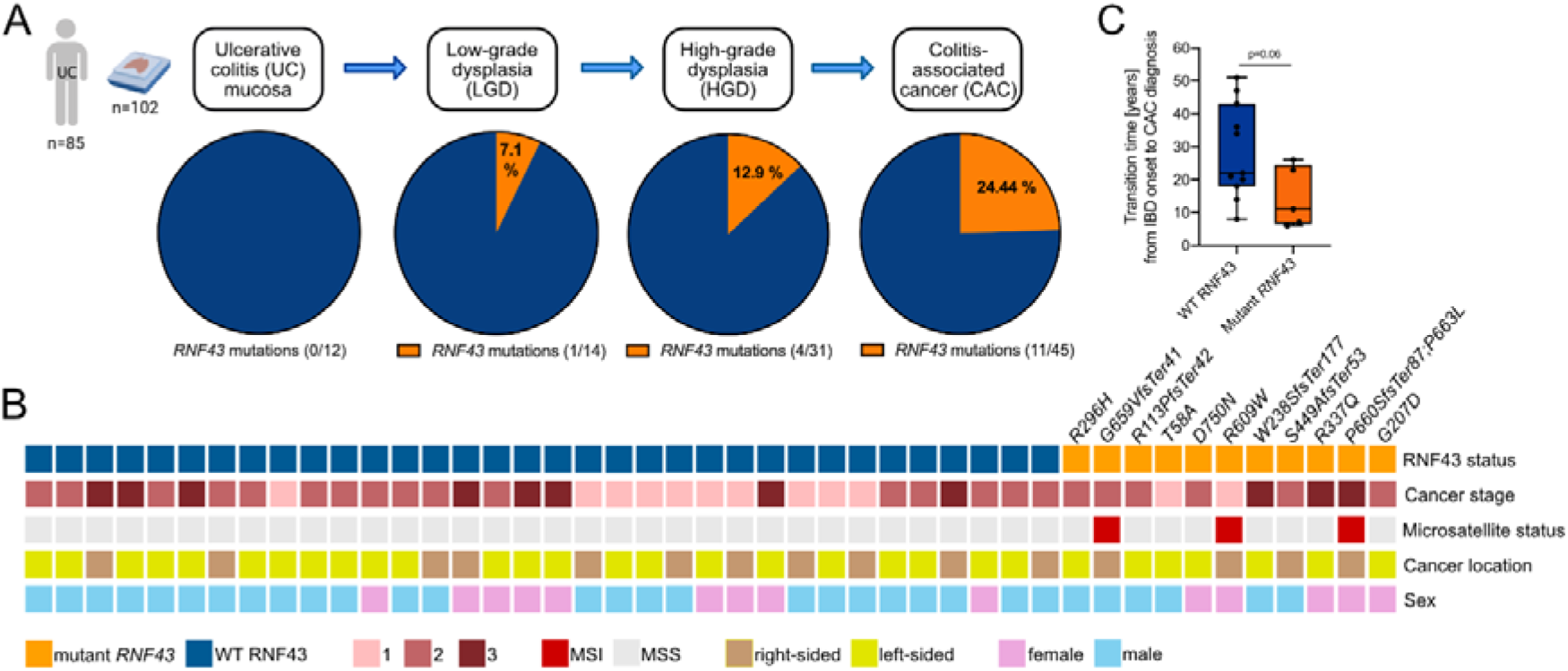
Mutations in *RNF43* are early events in the development of colitis-associated cancer. (A) Frequency of *RNF43* mutations in the inflamed colorectal tissue of UC patients (n = 85) with non-dysplastic mucosa (n = 12), LGD (n = 14), HGD (n = 31) and/or CAC (n = 45). (B) Clinical characteristics of CAC patients (n = 45), including RNF43 mutation and microsatellite status. (C) Transition time from UC onset until CAC diagnosis at cancer stages 1&2 in WT (n = 11) versus *RNF43* mutant patients (n = 5). Each symbol represents one patient. Bars indicate median. Unpaired T test was used.

### *Rnf43* mutations determine colitis severity in *Citrobacter rodentium-*infected mice and predispose to CAC development

To confirm whether mutations in *RNF43* drive the development of CAC, we established a mouse model mimicking human IBD in an altered *Rnf43* background by infecting RNF43^H292R/H295R^ and *Rnf43*^ΔEx8^ mice with the mouse pathogen *Citrobacter rodentium* (*C. rodentium*). We selected these two mouse lines based on our human findings (Figure 1B) and a human CAC study reporting that somatic mutations of *RNF43* are predominantly frameshift mutations located upstream of the RING domain (13), while infection with *C. rodentium* is a well-established model to study mucosal immunology and particularly IBD-like immune responses (16). As *C. rodentium* causes an acute, self-limiting infection, we analyzed mice at the peak of acute infection (12 days post infection (p.i.)) and in the late infection phase (28-36 days p.i.) (Figure 2A).

**Figure 2.**
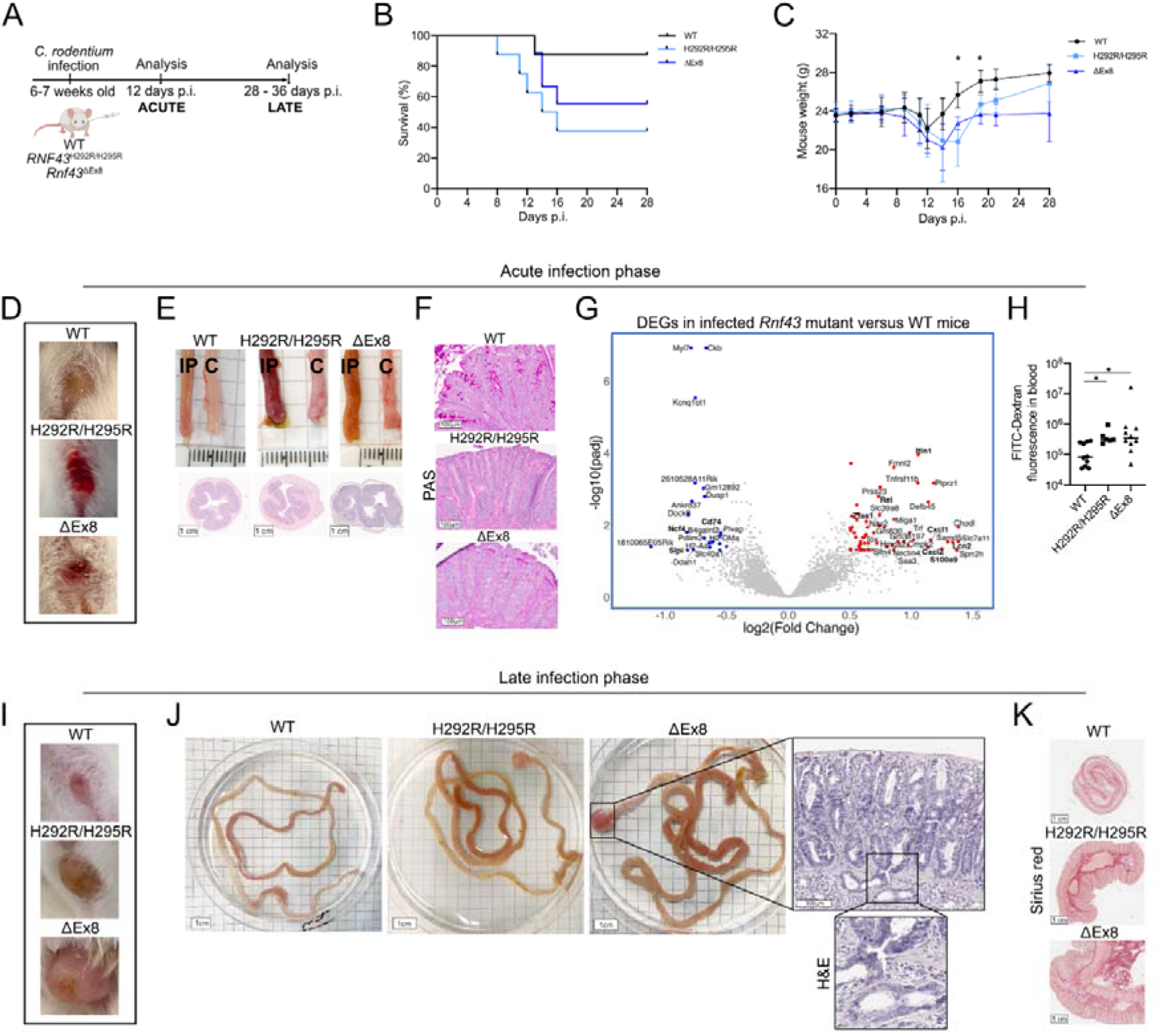
*Rnf43* mutations determine colitis severity in *Citrobacter rodentium-*infected mice and predispose to CAC development. (A) Experimental outline of *C. rodentium* infection. (B) Kaplan-Meier-curve of infected RNF43^H292R/H295R^ (n = 8), *Rnf43*^ΔEx8^ (n = 9) and WT (n = 8) mice. (C) Body weight (start weight 22.5 - 25 g) of *Rnf43* mutant versus WT mice during infection. (D-H) Acute colitis phenotype in *Rnf43* mutant versus WT mice. Representative appearance of (D) inflamed anorectal regions and (E) intestines (IP = intestine proximal, C = colon), with corresponding H&E-stained colorectal transections. (F) Representative PAS-stained colorectal sections displaying PAS^+^ goblet cells. (G) Volcano plot displaying the DEGs (upregulated: log_2_FC > 0.7, downregulated: log_2_FC <= -0.6) in the colorectal tissue of infected *Rnf43* mutant versus WT mice. (H) FITC-Dextran abundance measured in the blood. (I-K) Late colitis phenotype in *Rnf43* mutant versus WT mice. Representative appearance of (I) anorectal regions and (J) intestines with representative H&E-stained-colorectal prolapse showing invasive carcinoma development (zoom-in). (K) Representative Sirius red stained-colorectal sections displaying fibrosis. Each symbol represents one mouse. Bars denote median. Ordinary one-way ANOVA with Tukey’s multiple comparisons was used for normal distribution, otherwise Kruskal-Wallis-Test with Dunn’s multiple comparisons. *p < 0.05.

During the acute infection phase, only half of the *Rnf43* mutant mice survived (Figure 2B), showing substantial weight loss (Figure 2C), fecal abnormalities such as diarrhea, hematochezia and/or melena (Figure S1A), and severe anorectal inflammation (Figure 2D). Macroscopic inflammation signs such as intestinal reddening and thickening were more pronounced than in wild type (WT) mice (Figure 2E, upper panel), and reflected by increased intestinal diameters (Figure 2E, lower panel) and intestinal weight-to-length ratios (Figure S1B). Histological examination revealed pronounced erosion and loss of mucus-producing goblet cells (Figure 2F and S1C), which are histological hallmarks of UC (17). This exacerbated colitis phenotype in *Rnf43* mutant mice was also reflected at gene expression level, showing significantly upregulated inflammation markers (*Lcn2*, *S100A9*, *Rel, Cxcl1*, *Cxcl2)* and downregulated gut-repair- and barrier-proteins (*Cd74*, *Slpi*) (Figure 2G, Suppl. Table 1). A compromised barrier function is a key factor in IBD pathogenesis (18). In line with this, we observed a more severe intestinal barrier defect in *Rnf43* mutant mice, indicated by higher colorectal FITC-Dextran uptake into their blood (Figure 2H).

In the late infection phase, WT mice recovered, while *Rnf43* mutant mice developed colorectal prolapses (Figure 2I) and heavily thickened intestines (Figure 2J). In the prolapses, we observed adenomas and even invasive colorectal carcinomas (Figure 2J, zoom-in), which were embedded in massive fibrosis (Figure 2K).

These findings indicate that mutations in *Rnf43* are CAC driver mutations that confer susceptibility to aggravated and perpetuated colitis.

### *Rnf43* mutant mice show an impaired pathogen control and altered T cell response during *C. rodentium* infection

The observed colitis susceptibility in *Rnf43* mutant mice was reminiscent of IBD patients which show inappropriate immune responses directed against intestinal microorganisms (3). T cells are crucial in controlling *C. rodentium* (19), while aberrant T cell responses, and particularly pro-inflammatory CD4^+^ T cells, play an IBD-promoting role (20). Therefore, we examined *C. rodentium* colonization and T cell recruitment in our mouse model.

At the peak of infection, *Rnf43* mutant mice showed an increased *C. rodentium* load in the feces and a 100-fold higher colonization of the colon (Figures 3A and S2A), suggesting an impaired pathogen control. Simultaneously, significantly fewer CD3^+^ T cells infiltrated the colorectal epithelium (Figure 3B), in inverse manner to *C. rodentium* colonization (Figure 3C). CD3^+^ cells were also less abundant in the lamina propria (Figure 3B), but showed similar CD4^+^ T cell priming in terms of IFNγ- and IL-17A production as WT mice (Figure S2B), suggesting a compromised recruitment of T cells in *Rnf43* mutant mice.

**Figure 3.**
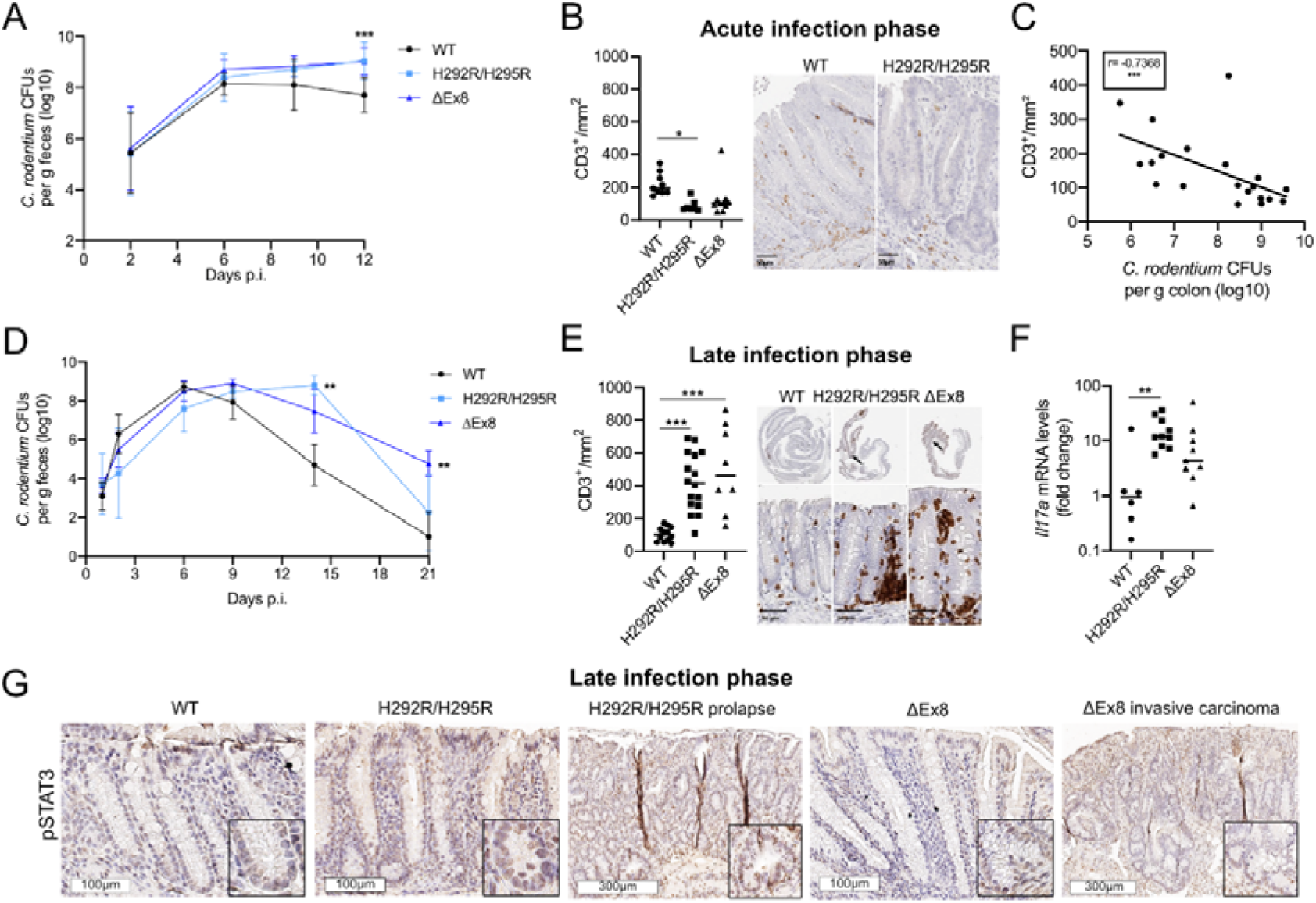
*Rnf43* mutant mice show an impaired pathogen control and altered T cell response during *Citrobacter rodentium* infection. (A-C) Acute infection phase in *Rnf43* mutant versus WT mice. (A) *C. rodentium* CFUs in the feces from RNF43^H292R/H295R^ (n = 7), *Rnf43*^ΔEx8^ (n = 10) and WT mice (n = 10) mice. (B) Intraepithelial CD3^+^ cells/mm^2^ colorectal tissue with representative CD3-stained sections. (C) Spearman correlation between colorectal CD3^+^ cells and *C. rodentium* CFUs. (D-G) Late infection phase in *Rnf43* mutant versus WT mice. (D) *C. rodentium* CFUs in the feces from RNF43^H292R/H295R^ (n = 5), *Rnf43*^ΔEx8^ (n = 6) and WT mice (n = 7) mice. (E) Intraepithelial CD3^+^ cells/mm^2^ colorectal tissue with representative CD3-stained sections. (F) Fold change in colorectal *Il17a* mRNA expression. (G) Representative pSTAT3-stained sections in prolapse, carcinoma and adjacent colorectal tissue of *Rnf43* mutant mice. Each symbol represents one mouse. Bars denote median. Ordinary one-way ANOVA with Tukey’s multiple comparisons was used for normal distribution, otherwise Kruskal-Wallis-Test with Dunn’s multiple comparisons. *p < 0.05, **p < 0.01, ***p < 0.001.

During the further course of infection, *C. rodentium* clearance was delayed in *Rnf43* mutant mice (Figure 3D) and associated with an exacerbated CD3^+^ cell recruitment to the mucosa (Figure 3E) that even exceeded the CD3^+^ numbers in WT mice at the peak of infection (Figure 3B). Upregulated colorectal *Il17a* gene expression together with a lowered *Foxp3/Il17a* ratio (Figures 3F and S2C) indicated a pro-inflammatory environment in the late infection phase in *Rnf43* mutant mice, which was confirmed by aberrant colorectal STAT3 signaling (Figure 3G). STAT3 governs Th17 cell differentiation and promotes CAC (21). In line with this, STAT3 activation gradually increased from adjacent colorectal tissue to prolapse and carcinoma lesions, and was particularly enriched in immune cells and malignant epithelial cells (Figure 3G).

Together, these observations indicate that mutated *Rnf43* is associated with an impaired pathogen control. Over time, the increased bacterial burden triggers an exacerbating Th17 response that appears to clear the pathogen, but potentially at the expense of increased tissue damage as *Rnf43* mutant mice eventually develop invasive colorectal carcinomas.

### Mutations in *Rnf43* alter intestinal goblet cell homeostasis and microbiota composition

To identify the factors that rendered *Rnf43* mutant mice more vulnerable to colitis, we characterized these mice under basal conditions. Already young (2 - 3 months old) *Rnf43* mutant mice exhibited reddened and thickened intestines with significantly elongated crypts (Figures 4A and S3A), suggesting an inflammatory phenotype that was not evident in other CRC models with aberrant WNT signaling like *Apc*^min^ mice (Figure S3B). We studied the colorectal gene expression in young *Rnf43* mutant mice for IBD-predisposing alterations and identified 160 differentially expressed genes (DEGs) (Suppl. Table 2). Genes involved in WNT signaling (*Axin2*, *Lgr5*, *Rnf43,* and *Pcdh17*) were highly upregulated (Figure 4B). Histological examination reflected this WNT signaling activation, as *Rnf43* mutant mice showed increased Ki67^+^ crypt proliferation (Figures 4C and S3C) and expanded OLFM4^+^ stem cell zones (Figure 4D) similar to *Apc*^min^ mice with high constitutive WNT activation. Also*, Rnf43* mutant colorectal stem cells gave rise to more and larger organoids (Figure S3D), further corroborating the increased stemness and proliferation upon epithelial *Rnf43* deficiency.

**Figure 4.**
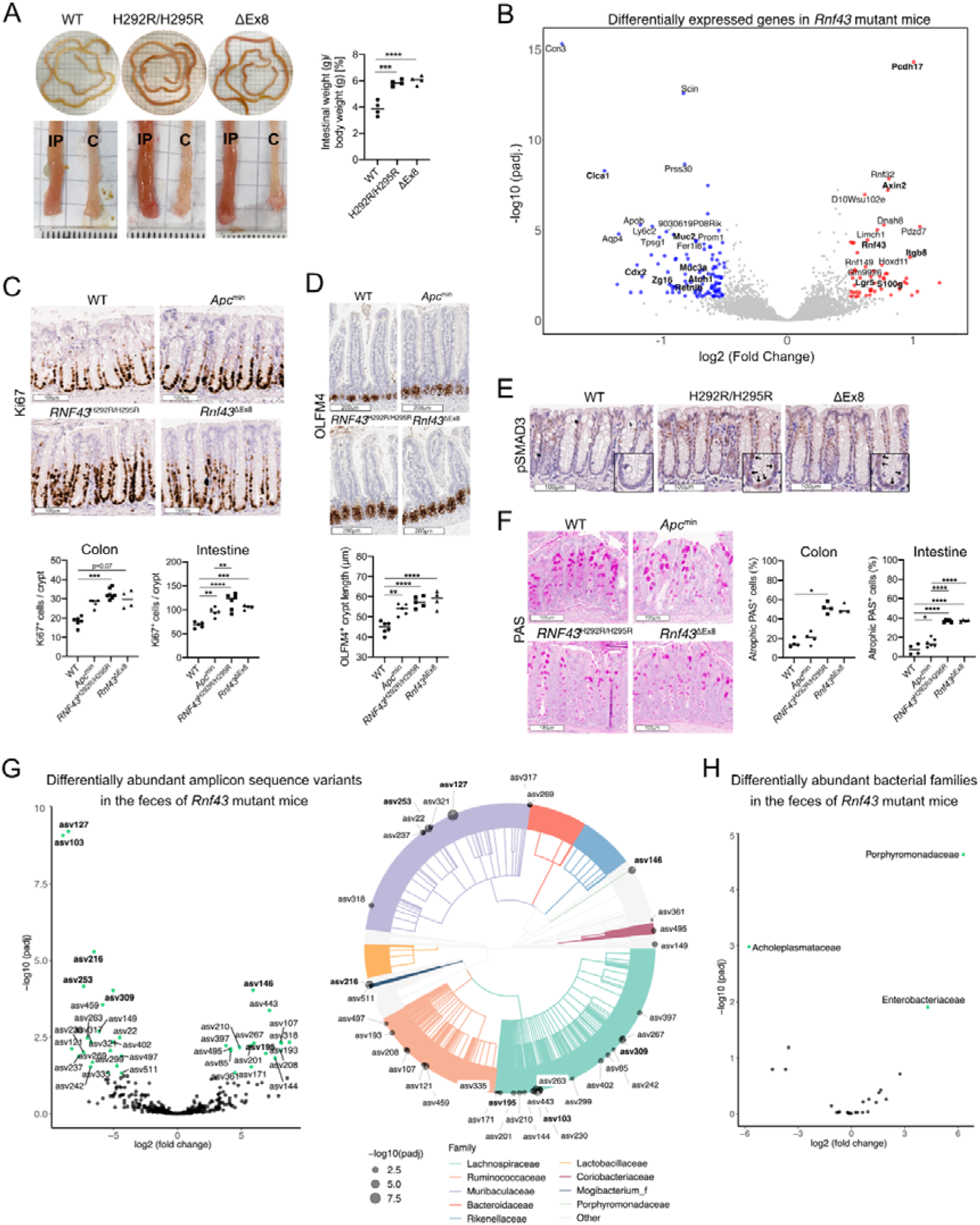
Mutations in *Rnf43* alter intestinal goblet cell homeostasis and microbiota composition. (A-H) Analysis of young (2-3 months old) RNF43^H292R/H295R^ and *Rnf43*^ΔEx8^ mice (FVB/N and/or C57BL/6 background). (A) Representative intestinal appearance (IP = intestine proximal, C = colon) and intestinal thickness quantified by intestinal weight per length. (B) Volcano plot displaying the DEGs (log2fc > 0.06, padj. < 0.002) in the colorectal tissue of *Rnf43* mutant versus WT mice. (C) Representative Ki67-stained colorectal sections with quantification of Ki67^+^ epithelial cells per crypt. (D) Representative OLFM4-stained intestines with length measurement of the OLFM4^+^ stem cell zone. (E) Representative pSMAD3-stained colorectal sections. (F) Representative PAS-stained colorectal sections with quantification of atrophic PAS^+^ goblet cells. 16S rRNA sequenced feces from *Rnf43* mutant versus WT mice showing (G) dASVs (log2fc > 0.3) with assignment to the bacteria family level and (H) differentially abundant families. Each symbol represents one mouse. Bars denote median. Ordinary one-way ANOVA with Tukey’s multiple comparisons was used for normal distribution, otherwise Kruskal-Wallis-Test with Dunn’s multiple comparisons. *p < 0.05, **p < 0.01, ***p < 0.001, ****p < 0.0001.

Furthermore, *Itgb8*, whose overexpression has been described as IBD risk factor (4), was highly upregulated in *Rnf43* mutant mice (Figure 4B). ITGB8 is a potent activator of latent TGF-β (22), which in turn plays a key role in IBD. Consistent with *Itgb8* overexpression, epithelial expression of pSMAD3, a downstream component of active TGF-β signaling, was increased in *Rnf43* mutant mice (Figure 4E).

Top downregulated genes (p_adj._ <= 0.0002, log_2_FC <= -0.7) included genes important for epithelial function (*Aqp4, Ffar2)* and epithelial differentiation (*Cdx2*), and particularly for goblet cell differentiation (*Atoh1*) and function (*Clca1*, *Muc2, Zg16, Retnlb (Relm*β*), Fer1l6, Muc3a, Ccn3, Scin, Tpsg1*) (Figure 4B), suggesting impairment of the epithelial mucus barrier function. Consistent with this, the number and particularly the size of mucus-secreting PAS^+^ goblet cells were reduced in *Rnf43* mutant mice (Figures 4F and S3E). This atrophic goblet cell phenotype was not present in *Apc*^min^ mice, suggesting it to be a WNT-independent effect of mutated *Rnf43*.

As mucus barrier and microbiota have an intricate relationship, we studied the fecal microbiota signatures in these young *Rnf43* mutant mice. Microbiota were strongly affected by the mutational status of *Rnf43* showing 37 differentially abundant amplicon sequence variants (dASVs), among which *Ruminococcus* (ASV 107) and *Parabacteroides goldsteinii* (ASV 146) were highly enriched (Figure 4G, Suppl. Tables 3-4). On the family level, we detected increased relative abundances of *Porphyromonadaceae* and *Enterobacteriaceae* (Figure 4H). *Enterobacteriaceae* play a crucial role in IBD pathogenesis (23), and also *Porphyromonadaceae* become enriched in IBD patients (24), and both correlate with CRC in humans and mice (25, 26), implicating the presence of IBD/CRC-promoting microbiota in young *Rnf43* mutant mice.

In summary, our data indicate that *Rnf43*-loss-of-function mutations activate epithelial WNT and TGF-β signaling, reduce goblet cell density, maturation and function, and cause dysbiosis.

### Mutations in *Rnf43* are sufficient to induce spontaneous intestinal inflammation with subsequent cancer development

To determine whether epithelial and goblet cell dysregulation together with dysbiosis initiate and drive spontaneous intestinal inflammation and inflammation-related tumorigenesis in *Rnf43* mutant mice over time, we characterized “aging” (9-13 months old) *Rnf43* mutant mice. These mice displayed further aggravated intestinal thickening and reddening (Figure 5A), and developed spontaneous tumors and rectal prolapses (Figure 5B), demonstrating that mutations in *Rnf43* are sufficient to induce severe intestinal pathological changes over time. The prolapse and tumor phenotype was detectable in RNF43 FVB/N, but also in two out of three C57BL/6 mice bearing mutant *Rnf43* (Figures 5C and S4A-S4B), thereby excluding a mouse strain background-driven effect.

**Figure 5.**
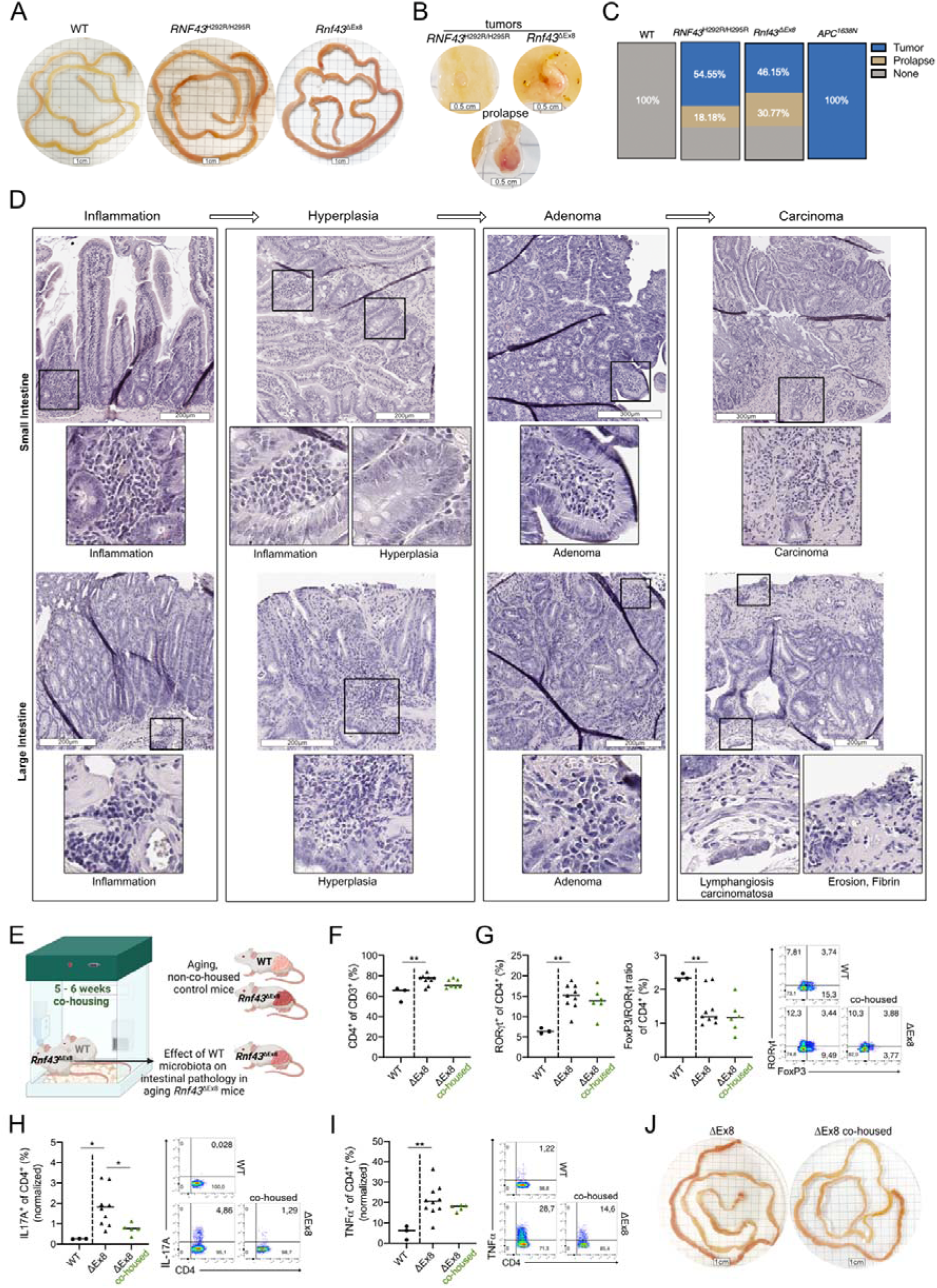
Mutations in *Rnf43* are sufficient to induce spontaneous intestinal inflammation with subsequent cancer development. (A-D) Macroscopic and histological phenotype of aging (9-13 months old) *Rnf43* mutant mice (FVB/N and/or C57BL/6 background). Representative (A) intestinal reddening, (B) tumors and rectal prolapses in *Rnf43* mutant mice. (C) Frequency of intestinal tumors and rectal prolapses in *Rnf43* mutant (*RNF43*^H292R/H295R^ n = 11, *Rnf43*^ΔEx8^ n = 13)*, Apc^1638N^* (n = 3) and WT (n = 7) mice. (D) Representative images of the inflammation-carcinoma-sequence observed in the small and large intestine of *Rnf43* mutant mice. (E-J) Small intestinal T cell phenotype in aging (9-11 months old) non-tumor-bearing *Rnf43*^ΔEx8^ mice compared to aging WT mice, and effects of WT microbiota on *Rnf43*-associated pathology. (E) Experimental outline of co-housing. Frequency and representative flow plots of (F) CD4^+^ T cells, (G) RORγt^+^ CD4^+^ subsets including FoxP3/RORγt ratio, (H) IL-17A^+^ CD4^+^ cells, and (I) TNFα^+^ CD4^+^ cells after stimulation with PMA/Iono (gated on: live, single cells, CD45^+^ and CD3^+^). (J) Representative intestinal appearance after co-housing. Each symbol represents one mouse. Bars denote median. Ordinary one-way ANOVA with Tukey’s multiple comparisons was used for normal distribution, otherwise Kruskal-Wallis-Test with Dunn’s multiple comparisons. *p < 0.05, **p < 0.01.

To estimate the contribution of aberrant WNT signaling to this phenotype, we compared *Rnf43* mutant mice to aging *Apc*^1638N^ mice, as those mice 1) have a WNT-driven phenotype and 2) reach such advanced age unlike *Apc*^min^ mice. *Apc*^1638N^ mice developed three times larger intestinal tumors than *Rnf43* mutant mice (Figure S4C), but did not show intestinal inflammation signs such as reddening or rectal prolapses (Figures S4A-4B), suggesting that inflammation is a major driver of tumorigenesis in *Rnf43* mutant mice. Supporting this assumption, *Rnf43* mutant mice displayed an UC-typical histopathology with mucosal erosion and fibrosis, and increased myeloid and lymphoid cell infiltration in close proximity to epithelial changes ranging from epithelial hyperplasia to adenomas and even invasive carcinomas (Figure 5D). These findings implicate that inactivated *RNF43* predisposes to inflammatory alterations over time in a WNT-independent manner.

We therefore hypothesized that the observed changes in the epithelial and mucus barrier render the intestinal epithelium of *Rnf43* mutant mice exposed to luminal microbiota, resulting in epithelial damage and mucosal inflammation. Thus, we studied whether aging *Rnf43* mutant mice (which showed microbiota alterations already at young age, Figures 4G-4H) displayed an IBD-like immune response before macroscopic tumor development, and whether it was influenced by *Rnf43*-microbiota. Therefore, we co-housed aging *Rnf43* mutant mice with young WT mice (Figure 5E). Their mutual fecal microbiota transfer allowed to investigate whether exposure to “healthy” microbiota from WT mice influenced the *Rnf43*-induced intestinal phenotype.

Aging *Rnf43* mutant mice harbored significantly higher proportions of CD4^+^ T cells than aging WT mice (Figures 5F and S4D). Those CD4^+^ cells showed a pro-inflammatory transcriptional signature, with increased subsets expressing RORγt, resulting in a declined FoxP3/RORγt ratio (Figure 5G). In line with this, IL-17A and TNFα producing CD4^+^ T cells, which are key mediators in IBD (27), were increased (Figures 5H and 5I). These results confirm the presence of an IBD-like T cell signature in aging *Rnf43* mutant mice.

6 weeks of co-housing with WT mice did not reduce the mucosal CD4^+^ T cell enrichment (Figure 5F), but significantly decreased the IL-17A-producing CD4^+^ T cells in co-housed *Rnf43* mutant mice (Figure 5H), suggesting that *RNF43*-associated microbiota contributes to pro-inflammatory Th17 priming. Intestinal reddening and rectal prolapses were also ameliorated in co-housed *Rnf43* mutant mice (Figure 5J). Intestinal thickening and spleen weight remained unchanged (Figures S4E-S4F), presumably due to the still existing goblet cell impairment and accumulated damage over time.

Together, these results underpin the significance of RNF43 in intestinal homeostasis, as inactivation of *Rnf43* alone is sufficient to initiate processes that drive the stepwise transformation of benign epithelial cells into (invasive) carcinomas. In comparison to WNT-driven tumorigenesis in *Apc* mutant mice, spontaneous tumor formation in *Rnf43* mutant mice is accompanied by a pronounced mucosal inflammation phenotype that seems to be linked to epithelial exposure to dysbiotic microbiota.

### Epithelial inactivation of *Rnf43* impairs goblet cell homeostasis and causes DNA-damaging and Th17/Th22-inducing microbiota

To dissect how microbiota alterations shaped by mutant *Rnf43* affect intestinal homeostasis and potentially goblet cells, we analyzed young WT mice after co-housing with aging *Rnf43* mutant mice (Figure 5E), and compared the findings to young WT and *Rnf43* mutant mice (Figure 6A).

**Figure 6.**
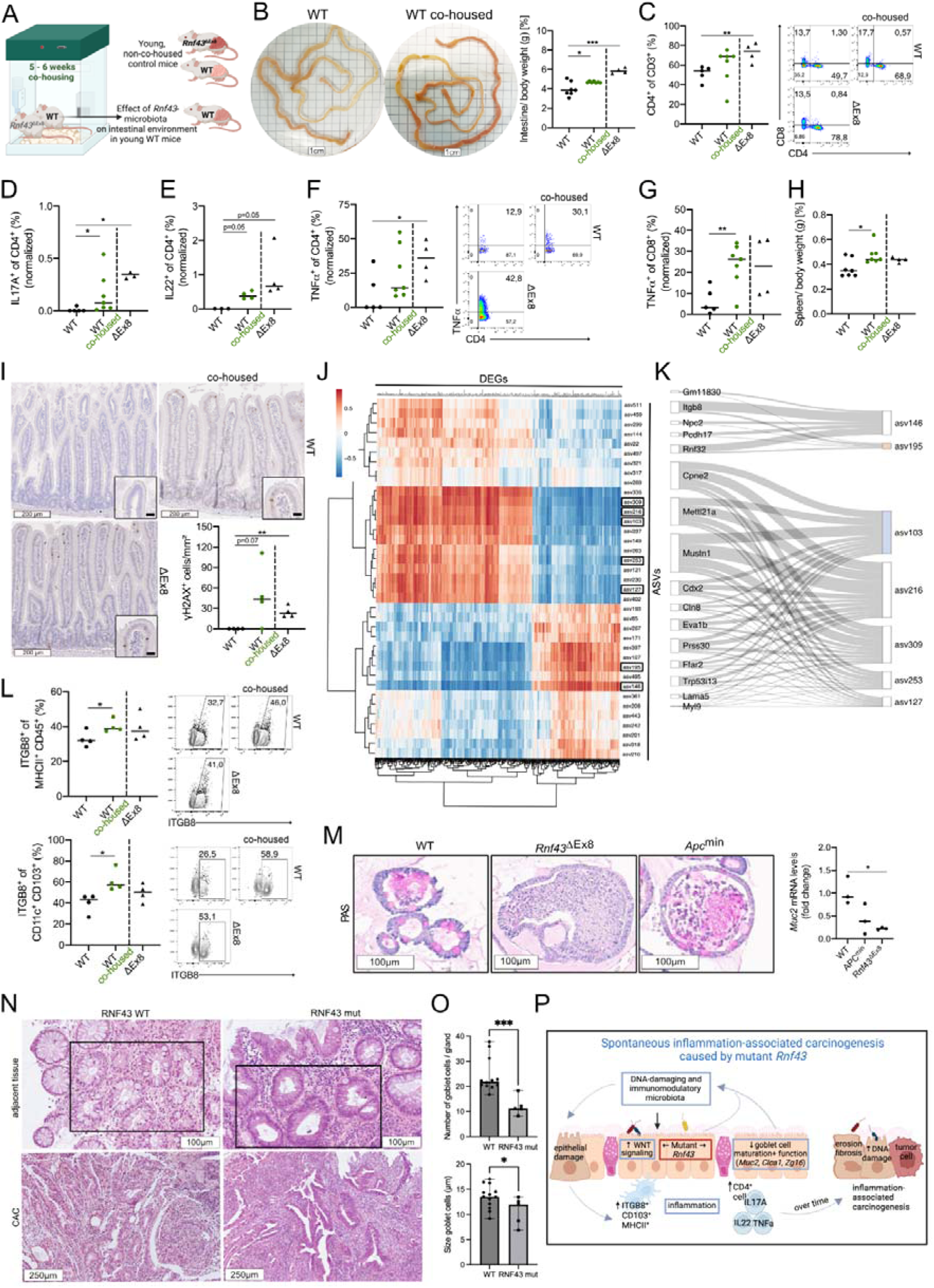
Epithelial inactivation of *Rnf43* impairs goblet cell homeostasis and causes DNA-damaging and Th17/Th22-inducing microbiota. (A-I) *Rnf43*^ΔEx8^-microbiota effects on intestinal homeostasis in young co-housed WT mice, compared to young WT and *Rnf43*^ΔEx8^ mice. (A) Experimental outline of co-housing. (B) Representative intestinal appearance and thickness quantified as intestinal weight to body weight ratio. (C) Frequency and representative flow plots of CD4^+^ T cells, which produce (D) IL-17A, (E) IL-22, (F) TNFα, and of (G) TNFα^+^ CD8^+^ cells after stimulation with PMA/Iono (gated on: live, single cells, CD45^+^ and CD3^+^). (H) Spleen weight after co-housing. (I) Representative γH2AX-stained small intestine with quantification of γH2AX^+^ epithelial cells/mm^2^. (J-M) Analysis of young (2-4 months old) *Rnf43* mutant mice. (J) Heatmap displaying estimated Kendall’s T correlation matrix between dASVs (log2fc > 0.3) and DEGs (log2fc > 0.5). (K) Sankey plot displaying the first sequential CCA step, gray links indicate positive DEG-ASV associations, missing links negative associations. (L) Frequency and representative flow plots of small intestinal ITGB8^+^ antigen presenting cells and DCs (gated on: live, single cells, CD45^+^ and MHCII^+^ and live, single cells, CD45^+^, MHCII^+^, CD4^-^, CD11c^+^ and CD103^+^). (M) Representative PAS^+^ stained-small intestinal organoids with *Muc2* mRNA levels. (N) Representative H&E-stained colorectal tissue specimens displaying RNF43 mutant and WT CAC and adjacent tissue areas (dysplastic crypts are framed), with quantification of goblet cell (O) numbers/gland (upper panel) and size (lower panel). (P) Graphical summary of the mechanisms by which mutant *Rnf43* causes spontaneous inflammation-associated colorectal cancer. Each symbol represents one mouse/patient. Bars denote median. Ordinary one-way ANOVA with Tukey’s multiple comparisons was used for normal distribution, otherwise Kruskal-Wallis-Test with Dunn’s multiple comparisons. *p < 0.05, **p < 0.01, ***p < 0.001.

The intestinal inflammation was partly phenocopied in young co-housed WT mice, which developed intestinal reddening and thickening (Figure 6B), and tended to have a higher proportion of CD4^+^ T cells compared to non-co-housed WT mice (Figure 6C). Functionally, these CD4^+^ cells showed a pro-inflammatory phenotype with increased IL-17A^+^, IL-22^+^ and TNFα^+^ subsets (Figures 6D-6F and S5A-S5B). Also, TNFα-producing CD8^+^ subsets (Figure 6G) and the spleen weight (Figure 6H) increased. Simultaneously, the DNA damage marker γH2AX increased in the epithelium of co-housed WT mice (Figures 6I). These findings demonstrate that the microbiota related to inactivated *RNF43* are a key driver of intestinal inflammation by inducing epithelial DNA damage together with pro-inflammatory T cell responses.

Furthermore, the comparison between non-co-housed WT and *Rnf43* mutant mice (Figure 6A) revealed the presence of a subacute inflammation in *Rnf43* mutant mice already at a young age, characterized by increased abundances of CD4^+^ T cells (Figure 6C) with elevated IL-17A, IL-22 and TNFα-producing subsets (Figures 6D-6F). These findings show that inflammation already exists months before tumors arise in *Rnf43* mutant mice, and confirm that the cancer cascade in *Rnf43* mutant mice is inflammation-driven, as in patients with IBD.

To establish a mechanistic link between microbiota alterations and molecular changes in *Rnf43* mutant mice, we computed a Kendall’s t correlation matrix between DEGs and dASVs (Figures 4B and 4G) in *Rnf43* mutant versus WT mice (Figure 6J), and identified a highly correlating subset that suggested intricate relationships between the top differentially abundant ASVs and a gene set comprising *Itgb8, Cdx2* and *Ffar2* (Figure 6K). ITGB8^+^ dendritic cells (DCs) increase in IBD patients upon inflammatory signals (28), and are important for Th17 differentiation (29). Increased abundances of ITGB8^+^ antigen presenting cells (MHCII^+^ CD45^+^) and particularly of ITGB8^+^ DCs were present in WT mice co-housed with *Rnf43* mutant mice (Figures 6L and S5C), providing a mechanism by which *Rnf43*-shaped microbiota can influence the adaptive immune cell compartment.

Of note, co-housing neither reduced goblet cell numbers and/or size in WT mice, nor did it normalize goblet cell alterations in *Rnf43* mutant mice (Figure S5D), implying that the goblet cell phenotype is not microbiota-driven but a consequence of the mutation. Importantly, *Rnf43* mutant stem cells generated organoids with fewer and atrophic goblet cells, and decreased *Muc2* gene expression (Figure 6M), thus confirming a cell-autonomous effect of inactivated *Rnf43* on goblet cell homeostasis. The impact of *RNF43* mutations on goblet cell numbers and size was also assessed in colorectal tissue samples from CAC patients. Since goblet cells are barely observed in CAC (Figure 6N), we analyzed the dysplastic tissue areas adjacent to CAC and observed that the number as well as the size of goblet cells was reduced in the colon of patients presenting mutated *RNF43* (Figures 6N and 6O).

In summary (Figure 6P), our findings demonstrate that *Rnf43* deficiency causes early spontaneous intestinal inflammation by impairing epithelial and particularly goblet cell homeostasis in a cell-intrinsic manner, and by causing dysbiosis. The dysbiosis is a key contributor to mucosal inflammation by inducing epithelial DNA damage and priming of TGF-ß-activating ITGB8^+^ DCs and pro-inflammatory (TNFα, IL-17A, IL-22) T cells. Over time, the continuous epithelial and goblet cell dysfunction with exposure to dysbiotic microbiota results in perpetuated and aggravated inflammation that drives accumulated epithelial damage with erosion, fibrosis and eventually invasive carcinoma development.

## DISCUSSION

Identification of the mutations and molecular mechanisms that drive carcinogenesis in IBD patients is important to discover early biomarkers for patient stratification, and to develop novel treatments that improve the poor prognosis characteristic of CAC. Mutations in *RNF43* were reported to be common in precancerous IBD lesions and CAC (13, 30), but whether and how these mutations contribute to inflammation and the progression from non-cancerous neoplasia to CAC remained unclear. We observed that *RNF43* mutations were already present in UC patients with low-grade dysplasia and increased stepwise during colitis-associated carcinogenesis (from 7.1 % in LGD and 12.9 % in HGD to 24.4 % in CAC). This early prevalence of mutant *RNF43* in UC patients, together with a shortened transition time from UC onset to CAC development in patients carrying mutations in *RNF43*, suggest that *RNF43* mutations are important CAC drivers. In line with this, we could demonstrate that mutations in *Rnf43* initiate and drive the biological processes that cause spontaneous intestinal inflammation and inflammation-driven cancers in mice.

Previous studies attributed the tumor suppressive function of RNF43 to WNT signaling regulation at different levels (31, 32). Intestinal effects of aberrant WNT signaling are very well-characterized in the *Apc* mutant CRC mouse model (33), and allowed us to discriminate WNT-mediated from WNT-independent intestinal alterations in *Rnf43* mutant mice. In young *Rnf43* mutant mice, we found upregulation of WNT target genes, and hyperplastic crypts with increase in stemness and proliferation similar to age-matched *Apc*^min^ mice, confirming the importance of RNF43 in regulating WNT signaling in the intestinal stem cell niche. However, we also identified a novel, WNT-independent role for RNF43 in intestinal homeostasis by regulating goblet cell morphology and physiology. We observed fewer and particularly atrophic intestinal goblet cells together with downregulation of goblet cell-specific genes (*Muc2, Atoh1, Clca1, Zg16, Retnlb, Fer1l6)* in *Rnf43* mutant mice, and a significant reduction in goblet cell density and size in colorectal tissue samples from CAC patients carrying mutated *RNF43*. A previous study reported a reduction in goblet cells in the presence of low *Rnf43*/*Znrf3* expression, and ascribed this reduction to aberrant WNT signaling in the epithelial stem cell niche, causing unrestricted proliferation and impaired differentiation (34).

However, the pronounced atrophic goblet cell phenotype in *Rnf43* mutant mice with significant *Muc2* downregulation in organoids was not present in *Apc*^min^ mice and organoids with highly aberrant WNT signaling activation, thereby linking *Rnf43* deficiency and goblet cell maturation in a cell-autonomous yet WNT-independent manner.

Secreted MUC2 serves as major structural component of the intestinal mucus barrier (35), which covers the intestinal epithelium like a dense blanket and stores antimicrobial peptides, thereby segregating gut microbiota from the epithelium (36). The significance of altered MUC2 expression in the development of mucosal inflammation is demonstrated by mice with *Muc2* knockout (*Muc2*^-/-^) and missense mutations (Winnie mice), and reflected by patients with active UC, who show reduced MUC2 production, less goblet cells and a thinned mucus layer (17, 37) that predisposes to facilitated epithelial-microbiota contact. As a consequence, young *Muc2*^-/-^ mice (complete goblet cell depletion) and Winnie mice (greatly decreased goblet cell number and size) develop intestinal inflammation that is characterized by mucosal thickening with hyperplastic crypts, epithelial erosion and increased T cell infiltration (38, 39). Later, these mice are prone to colitis-associated rectal prolapses, a complication found in UC (40), and carcinomas (12, 38). Young *Rnf43* mutant mice displayed a slower developing, but similar inflammation phenotype with reddened intestines and hyperplastic crypts, and increased proportions of pro-inflammatory (TNFα^+^, IL-17A^+^, IL-22^+^) CD4 T cells, which are key mediators in promoting IBD (27). Over time, *Rnf43* mutant mice frequently developed aggravated IBD-signs such as erosive and fibrotic alterations of the intestinal epithelium with mucosal myeloid and lymphoid cell infiltrates, as well as spontaneous rectal prolapses and tumors. Tumors ranked from adenomas to invasive carcinomas, and developed not only in the small but also in the large intestine. Invasive colorectal carcinomas are rarely seen in other CRC mouse models, making the *Rnf43* mutant mouse model a precious *in vivo* system to study the local colorectal processes involved in cancer development. Importantly, the striking inflammation phenotype shared by *Rnf43* mutant and *Muc2* mutant mice was not evident in *Apc* mutant mice with non-atrophic goblet cells, suggesting that impaired goblet cell maturation is a key contributor to the inflammation observed in *Rnf43* mutant mice. Thus, by displaying both features of *Apc* mutant mice (constitutively dysregulated epithelial WNT signaling activity) and of mice with genetic goblet cell alterations (goblet cell-mediated spontaneous mucosal inflammation), *Rnf43* mutant mice represent a unique colitis-associated cancer mouse model that allows to dissect the synergistic aspects and stages of spontaneous intestinal inflammation and inflammation-driven carcinogenesis beyond adenoma formation (inflammation -> hyperplasia -> adenoma -> carcinoma -> invasive carcinoma).

*C. rodentium* infection accelerated this spontaneous inflammation phenotype, and revealed that *Rnf43* mutant mice are predisposed to impaired pathogen control and pronounced barrier disruption, similar to *C. rodentium*-infected *Muc2-*deficient mice (41). In addition to MUC2, goblet cells produce and secrete the proteins CLCA1, ZG16 and RELMβ, which have bactericidal (42, 43), anti-tumorigenic (44) and immunomodulatory properties (45). Mice deficient for RELMβ suffer from impaired T cell recruitment after infection with *C. rodentium*, which then increasingly colonized and deeply penetrated the crypts (45) as observed in *Rnf43* mutant mice at the beginning of infection. Thus, downregulation of these goblet cell products in *Rnf43* mutant mice may additionally contribute to disruption of intestinal homeostasis.

Alterations in epithelial maturation and function in *Rnf43* mutant mice were accompanied by a characteristic fecal microbiota signature enriched in *Porphyromonadaceae* and *Enterobacteriaceae*, which is often seen in IBD patients (23, 24). Interestingly*, Porphyromonadaceae* (particularly *Parabacteroides goldsteinii,* which was among the most abundant bacteria in *Rnf43* mutant mice) and *Enterobacteriaceae* become enriched in CD2F1 mice upon subcutaneous injection of CRC cell lines (46), and increase in human adenocarcinomas (25), implying that their presence rises in cancerous environments. Changes in microbiota composition and mucus barrier mutually influence one another (8, 9). Fecal microbiota of *Rnf43* mutant mice did not induce goblet cell phenotype alterations in WT mice, and more importantly, *Rnf43* mutant intestinal organoids already displayed *Muc2* downregulation and reduction and atrophy of goblet cells *in vitro*, implying that the goblet cell phenotype observed is not microbiota-driven but a cell-intrinsic consequence of mutated *Rnf43*. The downstream effects of impaired goblet cell function on microbial communities (dysbiosis, inflammation) are very well-described (47). Considering this, it is tempting to speculate that the goblet cell alterations caused by epithelial RNF43 deficiency shape the gut microbiota.

Activation of STAT3 signaling, Th17/22 responses (48), production of reactive oxygen species (49), induction of DNA damage (50), and inhibition of tumor immunosurveillance (51) are reported mechanisms by which microbiota mediate inflammation, increase mutational burden, and accelerate tumorigenesis. In line with this, microbiota from *Rnf43* mutant mice induced intestinal inflammation in recipient WT mice characterized by intestinal reddening and thickening with increased IL-17A-producing CD4^+^ T cell abundances. Also, epithelial expression of the DNA-damage marker γH2AX as well as IL-22-producing CD4^+^ cells increased in these co-housed WT mice, indicating the pro-tumorigenic properties of the microbiota shaped by mutant *Rnf43.* IL-22 is secreted upon genotoxic stress and barrier impairment to support regeneration (52, 53). However, long-term triggered IL-22 secretion favors and perpetuates intestinal tumorigenesis (54), particularly through IL-22^+^ CD4^+^ T cell-mediated STAT3 activation in LGR5^+^ stem cells (55), and thereby fuels pro-tumorigenic pSTAT3 signaling, which indeed gradually increased in *Rnf43* mutant mice over time. Conversely, exposure to fecal WT microbiota ameliorated the intestinal inflammation phenotype in recipient *Rnf43* mutant mice, and underscored the benefits of fecal microbiota transfer over antibiotic treatment. Antibiotics have the drawback to promote colitis by depleting both pathogenic and healthy microbiota in patients (56), and to reduce goblet cells as well as increase CRC progression in *Apc*^min^ mice when long-term administered (57).

The most differentially abundant microbiota in young *Rnf43* mutant mice (*Parabacteroides goldsteinii* (ASV 146), *Ruminococcus* (ASV 107) and *Lachnospiraceae* (ASV 195, ASV 103)), correlated with the upregulation of *Itgb8*, and downregulation of *Cdx2* and *Ffar2*. ITGB8 activates latent TGF-β, and ITGB8^+^ DCs are central regulators of adaptive immunity by creating a TGF-β-rich milieu (58). ITGB8^+^ DCs increase in inflamed tissues of IBD patients (28), and were induced in WT mice upon co-housing, suggesting that the microbiota of *Rnf43* mutant mice may affect intestinal immunity through the ITGB8-TGF-β-axis. Numerous studies highlight the importance of dysregulated TGF-β signaling in IBD pathogenesis, when TGF-β turns from an immunosuppressive and protective into a harmful cytokine by promoting Th17 priming (59) and ECM-remodeling (60). Consistent with this, we found increased mucosal TGF-β signaling, Th17 priming and fibrosis in *Rnf43* mutant mice.

In summary, the microbiota shaped by mutant *Rnf43* cause epithelial DNA damage and prime pro-inflammatory and pro-tumorigenic (TNFα, IL-17, IL-22) T cell responses - with possibly systemic effects, as co-housed WT mice developed splenomegaly similar to *Rnf43* mutant mice. These microbiota-driven effects, together with the continuous epithelial and goblet cell barrier dysfunction and WNT dysregulation, cause the stepwise formation of inflammation-related cancers in *Rnf43* mutant mice.

Taken together, inactivation of *Rnf43* leads to spontaneous intestinal inflammation and inflammation-driven cancer. Screening of the mutational status of *RNF43* may improve early stratification and targeted treatment strategies for IBD patients at high CAC risk.

## ACKNOWLEDGEMENTS

We thank members of our laboratory ‘Chronic inflammation and carcinogenesis’ at TUM for technical support with special thanks to Karin Mink and Dr. Anna Brutau-Abia, and Dr. Karin Seidel and the MIH animal facility at TUM for their valuable support with animal welfare during *C. rodentium* experiments.

## AUTHOR CONTRIBUTIONS

A.D., R.M.L and M.G. conceived the study. A.D. designed, A.D. and A.R. performed and analyzed experiments. K.T., T.D., S.M., M.SK., D.S., R.O. and R.R. contributed to experiments. T.D., K.T. and R.M.L contributed to data analysis. A.R. and R.M.L contributed to data interpretation. M.ST. and R.O.A. performed computational analysis. C.L.M. supervised computational analysis. D.S. and X.L. contributed to computational analysis. T.G., K.S., W.S. and M.V. performed histopathological evaluation. M.V. provided human FFPE samples. M.V. and T.D. provided clinical information of CAC cohort. K.P.J. provided the *Apc*^1638N^ mouse model. A.D. and R.M.L. wrote the article. M.G. and R.M.L. acquired the funding. All authors read and reviewed the article.

A.D. held the “Medical Life Science and Technology” PhD Scholarship granted by the TUM Medical Graduate School. T.D. was supported by the Forschungskommission Klinikum Bayreuth (#50018). M.ST. held the "Munich School for Data Science - MUDS" PhD Studentship by the Helmholtz Association.

## DATA AVAILABILITY

Raw RNA sequencing and 16S rRNA sequencing data will be made available under the BioProject ID PRJNA951425.

## ABBREVIATIONS USED

dASVs: (differentially abundant amplicon sequence variants),
CAC: (colitis-associated cancer), canonical correlation analysis
(CCA): *C. rodentium* (*Citrobacter rodentium*),
CRC: (colorectal cancer),
DCs: (dendritic cells),
DEGs: (differentially expressed genes),
ECM: (extracellular matrix), formalin-fixed and paraffin-embedded
(FFPE): Hematoxylin & Eosin (H&E),
HGD: (high-grade dysplasia),
IBD: (inflammatory bowel disease),
LGD: (low-grade dysplasia),
(PAS): Periodic acid staining,
(p.i.): post-infection,
RNF43: (Ring finger protein 43),
UC: (ulcerative colitis),
WT: (wild type).

## MATERIAL AND METHODS

### Human cohort

102 FFPE colorectal tissue samples from 85 UC patients were categorized into four pathological stages of colitis-associated carcinogenesis (Figure 1). Inflamed, non-dysplastic mucosa samples (n = 12) from this cohort served as control. The study was approved by the ethics committee of the University Bayreuth, Germany (Az.O1305/1-GB).

**Figure 1:**
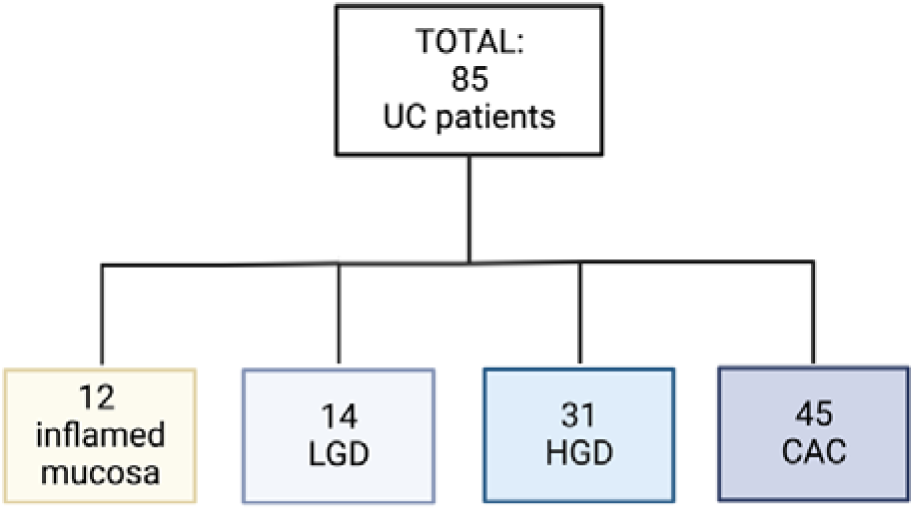
Overview of the 102 UC patient samples included in this study. From three patients both CAC and HGD samples were analyzed, from one patient both CAC and LGD samples, and from one patient both HGD and LGD samples. UC = ulcerative colitis, LGD = low-grade dysplasia, HGD = high-grade dysplasia, CAC = colitis-associated cancer.

### Exome sequencing

Genomic DNA was isolated from the 102 human FFPE colorectal tissue samples using the Maxwell^®^ RSC Blood DNA Kit (Promega) on a Maxwell^®^ 16 MDx (Promega). RNF43 exome sequencing was performed using a *RNF43* Custom Panel (Illumina). Paired-end sequencing was carried out on a MiSeq instrument (Illumina, Inc.). Analysis was performed with the DNA Amplicon App v2.1.1 on BaseSpace Sequence Hub Professional and Variant Interpreter v2.11.1.1 (Illumina). Single nucleotide polymorphisms and INDELS with a quality score of > 90, an allelic frequence of > 5 % and a population frequency < 1% were considered as somatic variants.

### Microsatellite-status

Microsatellite-instability (MSI) was determined in all human UC samples by fragment length analysis of six loci (BAT-25, BAT-26, NR-21, NR-22, NR-24, NR-27) with quasimonomorphic mononucleotide repeats (Table 1). After MSI multiplex PCR and capillary electrophoresis, samples were analyzed on a SeqStudio^TM^ Genetic Analyzer v1.1.4 (Thermo Fisher) with SeqStudioTM Plate Manager v1.0 (Thermo Fisher), and with GeneMapper^®^ v5 (Thermo Fisher). The same validated microsatellite-stable patient served as control, and MSI was defined when ≥ 2 instable markers were present in all six analyzed loci.

**Table 1:**
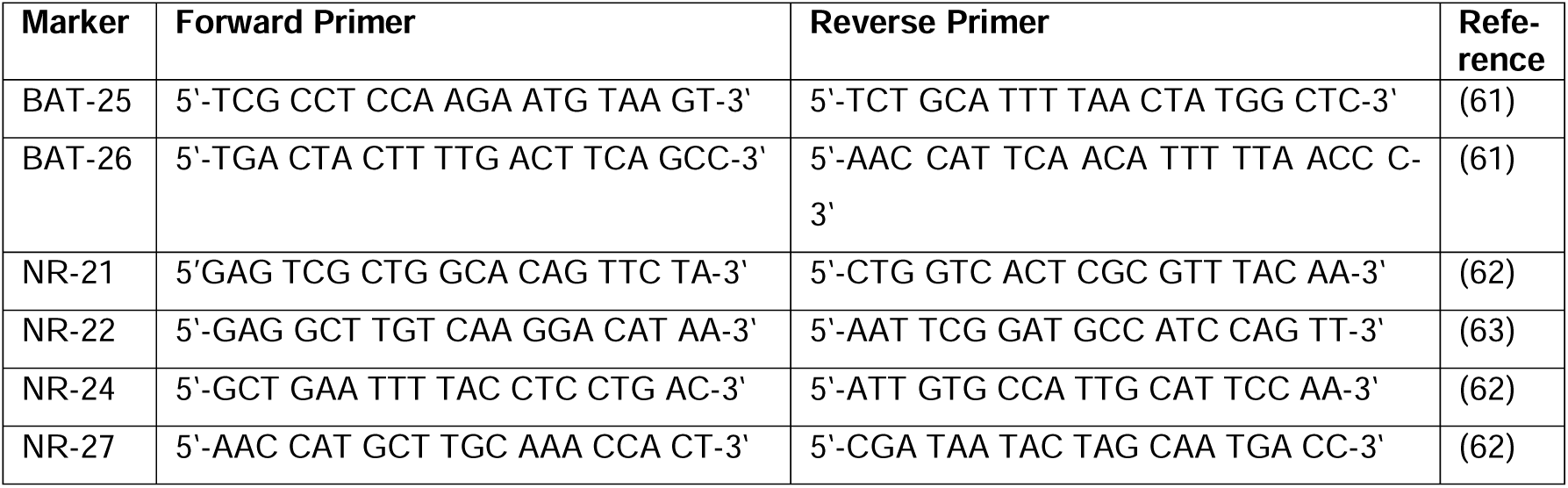
Panel of microsatellite markers.

### Mouse models

RNF43^H292R/H295R^ mutant mice (two point mutations in the RING domain) and *Rnf43*^ΔEx8^ mice (devoid of the RING domain) were used in the FVB/N (64) and C57BL/6 background. As WNT controls, *Apc*^+/min^ mice (C57BL/6J-*Apc^Min^*/J; RRID:IMSR JAX:002020) and *Apc*^+/1638N^ mice (provided by Prof. Klaus-Peter Janssen, TUM, Germany) were used. Mice were bred under specific pathogen-free conditions, and received a standard diet and water ad libitum. All animal experiments were conducted in accordance with the European guidelines for the care and use of laboratory animals after approval by the Bavarian Government (Regierung von Oberbayern, Az.55.2-1-54-2532-127-2017).

### Citrobacter rodentium culture and infection

GFP-labeled, kanamycin-resistant *Citrobacter rodentium* (kindly provided by Prof. Siegfried Hapfelmeier, University of Bern, Switzerland) (65), was cultured in a microaerophilic atmosphere, first on MacConkey agar II plates (BD) and then in Luria Broth, both supplemented with kanamycin (C. Roth). Once the exponential bacterial growth phase was reached, 1 × 10^8^ *C. rodentium* were administered to 6 – 7 weeks old mice per oral gavage.

### Citrobacter rodentium colony forming units

*C. rodentium* colony forming units were determined based on serially diluted and plated supernatants of feces and/or colorectal tissue homogenates (0.1 g feces or tissue homogenized in 1 ml PBS) as follows:

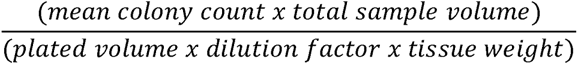

### FITC-Dextran assay

150 µl of 80 mg/ml FITC-Dextran (4kDa, Sigma) were orally administered to fasted mice. 4 - 6 hours later, FITC fluorescence intensity in the blood was measured (485 nm excitation, 535 nm emission) using the SoftMax Pro 7.1 program on a spectrofluorometer (SpectraMax i3X), and normalized to PBS blanks and blood fluorescence baseline levels.

### Histology and Immunohistochemistry

Formalin-fixed (in 10 % neutral-buffered formalin for 48 hours) and paraffin-embedded (FFPE) tissues were cut into 4 µm thick sections, deparaffinized and rehydrated for staining. For immunohistochemical staining, peroxidase treatment was followed by antigen retrieval with sodium citrate (10 mM, pH 6) or EDTA (1 mM, pH 8) buffer, and primary antibodies were incubated overnight at 4°C (Table 2). Horseradish peroxidase-conjugated secondary antibodies (Promega) and diaminobenzidine (Cell Signaling) were applied to amplify and detect the signal. Periodic acid Schiff (PAS) reaction was conducted for 5 min in 1 % periodic acid (C. Roth) and 15 min in Schiff’s reagent (C. Roth), Sirius Red staining for 1 hour in picric Sirius Red (Sigma Aldrich). Stained slides were scanned by the Aperio AT2 brightfield slide scanner (Leica Biosystems). Independent researchers evaluated the intestinal sections blindly by counting positive cells per area or by measuring the length of a functional unit (intestinal crypt/villus unit or colorectal crypt) using Aperio ImageScope (v.12.4.0.7018, Leica BioSystems). Goblet cells were considered atrophic with a cell size <= epithelial cell nuclei.

**Table 2.** Antibodies used for immunohistochemical analysis.

| Target | Clone | Origin/Target | Antigen retrieval | Dilution | Company |
| --- | --- | --- | --- | --- | --- |
| OLMF4 | D6Y5A | rabbit | sodium citrate (10mM, pH6) | 1:400 | Cell Signaling (#39141) |
| Ki67 | D3B5 | rabbit | sodium citrate (10mM, pH6) | 1:400 | Cell Signaling (#12202) |
| CD3 | SP7 | rabbit | sodium citrate (10mM, pH6) | 1:150 | ThermoFisher (#RM-9107) |
| pSMAD3 | EP823Y | rabbit | sodium citrate (10mM, pH6) | 1:250 | Abcam (#ab52903) |
| pSTAT3 | D3A7 | rabbit | EDTA (1mM, pH8) | 1:200 | Cell Signaling (#9145) |
| γH2AX | EP854(2)Y | rabbit | sodium citrate (10mM, pH6) | 1:3000 | Abcam (#Ab81299) |
| HRP-conjugated anti-rabbit |  | anti-rabbit |  | 1:200 | Promega (#W4011) |

Histopathological evaluation and tumor grading of mouse intestines and tumors was performed based on the current, published nomenclature and diagnostic criteria for murine gastrointestinal lesions (66).

### Generation and culture of small and large intestinal organoids

2-4 mm minced intestinal tissue pieces were washed and incubated (4 °C, 15 minutes, for 3 times) in EDTA (large intestine in 30 mM, small intestine in 2.5 mM). The filtered crypts were seeded in 40 µl basement membrane matrix Matrigel (Thermo Fisher) and cultured in 500 µl organoid growth medium (Tables 3 and 4). L-Wnt3a, R-Spondin 3 and Noggin (L-WRN) conditioned medium was obtained by culturing L-WRN producing cells (ATCC CRL-3276).

**Table 3.** Organoid growth medium components.

|  |
| --- |
| Single crypt medium |
| 50 % DMEM/F12 |
| 50 % L-WRN (L-Wnt3a, R-Spondin 3 and Noggin) |
| 1% Glutamax |
| 1 % Hepes |
| 1 % Penicillin/Streptomycin |

**Table 4.** Supplements for the organoid growth medium.

| Supplement | Dilution | Catalog # | Company |
| --- | --- | --- | --- |
| EGF | 1:1000 | PMG8043 | Gibco |
| B27 | 1:50 | 17504-044 | Gibco |
| N2 | 1:100 | 17502-048 | Invitrogen |
| N-Acetylcystein | 1:400 | A9165 | Sigma-Aldrich |
| Y-27632 | 1:1000 | 72304 | Stemcell Technologies |
| Cell recovery solution | 500 µl/well | 354253 | Corning |
| Matrigel Basement Membrane Matrix, growth factor reduced (GFR) | 40 µl/well | 47743-720 | ThermoFisher |

### Downstream analysis of intestinal organoids

The organoid-matrigel dome was dissolved in cell recovery solution (Corning) for subsequent RNA isolation or incubated in 4 % PFA for histological staining and analysis. Organoid growth was assessed taking overview images over a period of six days (EVOS FL Auto 2 imaging system, Thermo Fisher Scientific), and the mean size of up to forty organoids per day and mouse was determined using ImageJ software.

### Lamina propria leukocyte isolation and flow cytometry

Intestinal tissue was incubated in 30 mM EDTA (Invitrogen), and digested with 0.5 mg/mL collagenase from Clostridium histolyticum Type IV (Sigma Aldrich) and 10 μg / mL DNase I (Applichem). Filtered single cell suspensions were blocked (anti-mouse CD16 / CD32) and stained for live/dead discrimination (Zombie Aqua, BioLegend, or EMA, Thermo Fisher) prior to staining with fluorophore-conjugated antibodies (30 min) (Table 5). FoxP3 Transcription Factor Staining Buffer Set (eBioscience) and intracellular cytokine staining kit (BD Biosciences) were used according to manufacturer’s instructions. Prior to intracellular staining, cells were stimulated with Phorbol myristate acetate/Ionomycin (PMA/Iono, Sigma-Aldrich) and incubated with protein transport inhibitor Golgi Plug (BD Biosciences) at 37° C for 4 hours. Cells were acquired on a 4 laser CytoFlex S (Beckman Coulter) and analyzed using FlowJo software (V10.8.0).

**Table 5.** Antibodies and kits used for flow cytometry.

| **Epitope** | **Fluoro-chrome** | **Clone** | **Dilution/**<br>**Concentration** | **Catalog #** | **Company** |
| --- | --- | --- | --- | --- | --- |
| anti-mouse CD16/CD32 | - | 93 | 1:500 | 14-0161 | eBioscience |
| Zombie Aqua Fixable Viability kit | - | - | 1:1000 | 423102 | BioLegend |
| Foxp3 / Transcription Factor Staining Buffer Set | - | - | - | 00-5523 | Thermo Fisher |
| BD Cytofix/Cytoperm™ Fixation/Permeabilization Kit | - | - | - | 554714 | BD Biosciences |
| BD GolgiPlug™ Protein Transport Inhibitor | - | - | 1:1000 | 555029 | BD Biosciences |
| Phorbol myristate acetate (PMA) | - | - | 200 ng/mL | P1585 | Sigma-Aldrich |
| Ionomycin (Iono) | - | - | 500 ng/mL | I3909 | Sigma-Aldrich |
| anti-mouse CD45 | AlexaFluor 700 | 30-F11 | 1:400 | 103128 | BioLegend |
| anti-mouse CD3ε | FITC | 500A2 | 1:200 | 152304 | BioLegend |
| anti-mouse CD4 | BV605 | RM4-5 | 1:250 | 100548 | BioLegend |
| anti-mouse CD4 | eFluor450 | RM4-5 | 1:250 | 48-0042 | eBioscience |
| anti-mouse CD4 | PE Dazzle 594 | GK1.5 | 1:250 | 100455 | BioLegend |
| anti-mouse CD8a | APC-H7 | 53.6-7 | 1:250 | 560182 | BD Biosciences |
| anti-mouse FoxP3 | eFluor450 | FJKL-16s | 1:200 | 45-5773 | eBioscience |
| anti-mouse FoxP3 | PerCP Cy5.5 | FJKL-16s | 1:200 | 45-5773 | eBioscience |
| anti-mouse RORγt | PE | AFKJS-9 | 1:100 | 12-6988 | eBioscience |
| anti-mouse IL-17A | APC | eBio17B7 | 1:150 | 17-7177 | eBioscience |
| anti-mouse TNFα | PE Dazzle 594 | MP6-XT22 | 1:200 | 506346 | BioLegend |
| anti-mouse IL-22 | PE | Poly5164 | 1:150 | 516404 | Biolegend |
| anti-mouse MHCII | PE | M5/114.15.2 | 1:1000 | 12-5321 | eBioscience |
| anti-mouse CD103 | BV605 | 2E7 | 1:100 | 121433 | BioLegend |
| anti-mouse CD11b | APC Cy7 | M1/70 | 1:200 | 101225 | BioLegend |
| anti-mouse CD11c | APC | N418 | 1:200 | 17-0114 | eBioscience |
| ITGB8 | unconjugated |  | 1:50 | PA5-97052 | Thermo Fisher |
| Chicken anti-rabbit | Alexa Fluor 488 | Polyclonal | 1:200 | FAB8165G | R&D Systems |

### RNA isolation, cDNA synthesis and quantitative real-time-PCR

Intestinal tissue was homogenized with a Precellys^®^ 24 homogenizer (Avantor). RNA was isolated with the Maxwell 48 RSC simply RNA Tissue Kit (Promega) on a Maxwell RSC Instrument (Promega). Moloney Murine Leukemia Virus Reverse Transcriptase RNase H-Point Mutant (Promega) was used for cDNA synthesis, and GoTaq qPCR Mastermix (Promega) for quantitative real-time PCR on a CFX384 instrument (Bio-Rad). Conditions applied included 40 cycles of amplification, 15 sec denaturation at 95 °C, 30 sec annealing and amplification at 60 °C. CT values of target genes (Table 6) were normalized to *Gapdh* and to WT mice (2^-ΔΔCT^ method).

**Table 6.**
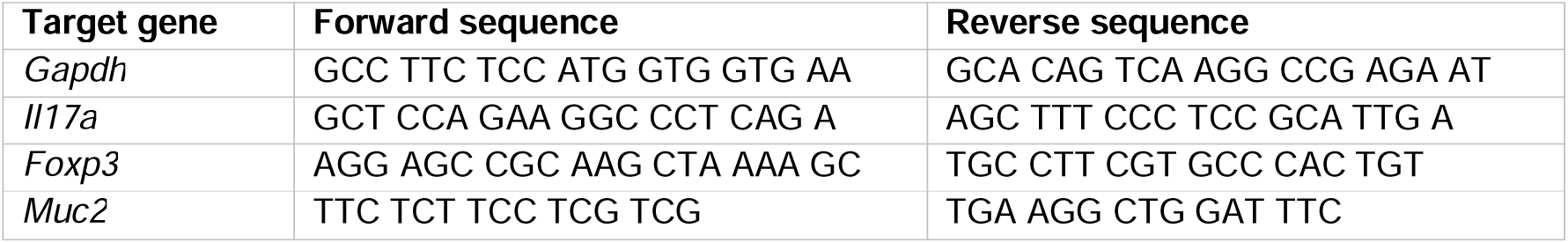
Primer sequences used for qPCR.

### Bulk RNA sequencing and differential gene expression analysis

After testing the RNA integrity via formation of clear 28S and 18S rRNA bands in gel electrophoresis, library preparation for bulk poly(A)-RNA sequencing was done as described previously (67), except for exchanged P5 and P7 sites to allow sequencing of the cDNA in read1 and of the barcodes and unique molecular identifiers (UMIs) in read2 to achieve a better cluster recognition. The library was sequenced on a NextSeq 500 (Illumina) (59 cycles in read1 and 16 cycles in read2). Data was processed using Drop-seq v1.0 to generate sample- and gene-wise UMI tables (68). The reference genome (GRCm38) was used for alignment. GENCODE version M25 was used for transcript and gene definitions.

Differential gene expression analysis was performed with DESeq2 after removing genes with less than 10 reads. *Rnf43* mutant (*Rnf43*^ΔEx8^ and RNF43^H292R/H295R^) mice were compared to wild type mice while controlling for co-housing, pregnancy, and experiment batch. Genes with an adjusted p-value of 0.05 were considered as differentially expressed.

### 16S rRNA sequencing and differential amplicon sequence variant analysis

For bacterial DNA extraction, feces was homogenized with a Precellys® 24 homogenizer (Avantor), and a modified Phenol-Chloroform-Isoamyl alcohol extraction (adapted from 69) and ethanol-precipitation method (adapted from 70) was applied. The V3/V4 region of the 16S rRNA gene was amplified using 341F-ovh and 785R-ovh primers (71). Paired-indexed (Nextera XT indices, Illumina) and magnetic bead-purified (AMPureXP, Beckman Coulter) samples were sequenced for 600 cycles using the MiSeq reagent kit v3 (Illumina). After trimming with Cutadapt (72), samples were processed with DADA2 v1.18 (73), and assignment of amplicons to amplicon sequence variants (ASVs) was performed using EzBiocloud bacterial database (74) in Mothur with classify.seqs command (Bayesian cut-off of 80 %).

Differentially abundant ASVs (dASVs) between *Rnf43* mutant (*Rnf43*^ΔEx8^ and RNF43^H292R/H295R^) and WT mice were determined on longitudinal sets of ASVs (feces sampled on day 0, 2, 6, and 12) using LinDA v.0.1.0. (75), while controlling for experiment batch, co-caging, co-housing, pregnancy of one mouse, time point of analysis and mouse ID. All ASVs with a Benjamini/Hochberg adjusted p-value <= 0.05 were considered differentially abundant.

### Computational transcriptome-microbiome-associations

For microbiome-transcriptome-associations, a sparse canonical correlation analysis (CCA) was performed for the longitudinally dASVs (upregulated: padj. < = 0.05 and log_2_FC > 0.3; downregulated: padj. < = 0.05 and log_2_FC < -0.3) and DEGs (upregulated: padj. < 0.05 and log_2_FC > 0.5; downregulated: padj. < 0.05 and log_2_FC < -0.5) between *Rnf43* mutant and WT mice using mixedCCA v.6.2.1. (76). Genes were variance stabilized and ASVs were transformed (modified clr) prior to analysis. It was assumed that genes follow a continuous distribution, ASVs a truncated (zero-inflated) continuous distribution. For variable selection in mixedCCA BICtype = 2 was used.

### Statistical analysis

Depending on Gaussian distribution, statistical significance between two groups was determined with unpaired student’s t-test or Mann-Whitney-U test, and for analysis among more than two groups one-way ANOVA with Tukey’s multiple-comparisons test or Kruskal-Wallis-test with Dunn’s multiple-comparisons test. P values < 0.05 were considered significant. Exact p values are stated when relevant. Statistical analysis was carried out using Prism 8 (GraphPad Software). Graphical illustrations were created with BioRender.com, and Affinity Photo and Affinity Designer (Serif) were used for figure design.

---

**Suppl. Figure 1.**
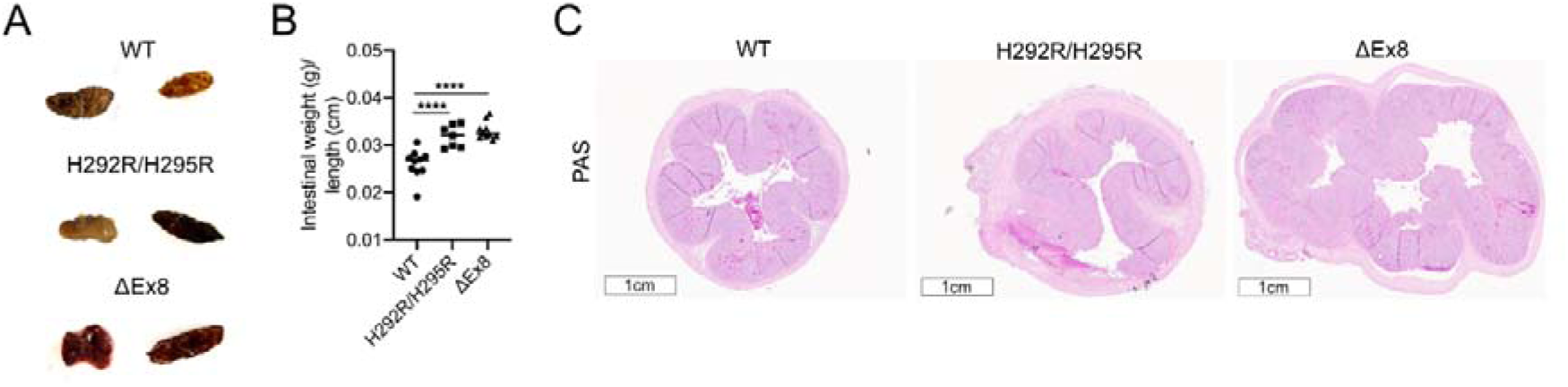
Mutations in *Rnf43* determine colitis severity in *Citrobacter rodentium* infected mice and predispose to CAC development. (A-C) Acute *C. rodentium* colitis signs in *Rnf43* mutant versus WT mice. (A) Mucoid feces (middle panel), hematochezia (lower panel) and melena (middle and lower panel) in infected *Rnf43* mutant mice. (B) Intestinal thickness measured by calculating the intestinal weight over intestinal length. (C) Representative PAS-stained colorectal transections displaying PAS^+^ goblet cells. Each symbol represents one mouse. Bars indicate median. Ordinary one-way ANOVA with Tukey’s multiple comparisons correction was used for normal distribution, otherwise Kruskal-Wallis-Test with Dunn’s multiple comparisons correction, unless indicated differently. ****p < 0.0001.

**Suppl. Figure 2.**
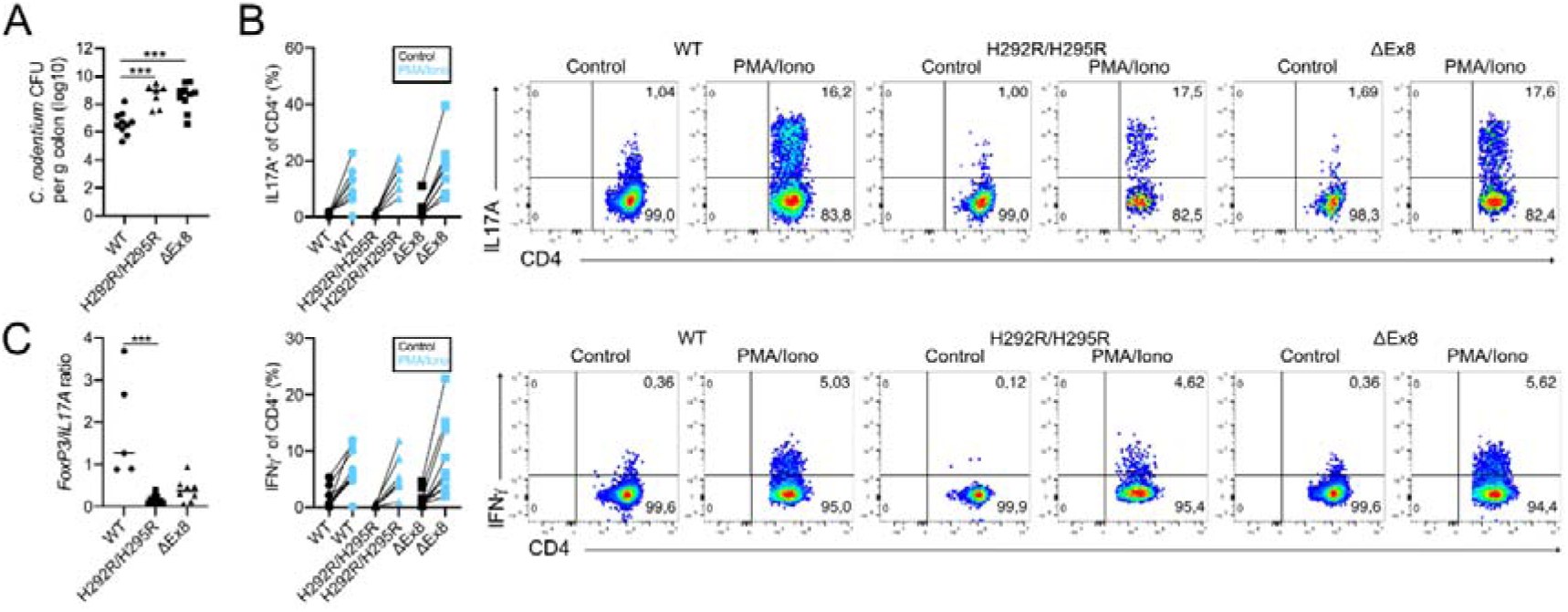
*Rnf43* mutant mice show an impaired pathogen control and altered T cell response during *C. rodentium* infection. (A) *C. rodentium* CFUs in colorectal tissue at the peak of infection (12 days p.i.). (B) Frequency and representative flow plots of CD4^+^ colorectal T cells producing IL-17A (upper panel) and IFNγ (lower panel) after stimulation with PMA/Iono in the acute infection phase. Cells were pre-gated on live, single cells, CD45^+^, CD3^+^, CD4^+^. (C) Colorectal *Il-17a* and *Foxp3* mRNA expression ratio in the late infection phase. Each symbol represents one mouse. Bars indicate median. Ordinary one-way ANOVA with Tukey’s multiple comparisons correction was used for normal distribution, otherwise Kruskal-Wallis-Test with Dunn’s multiple comparisons correction, unless indicated differently. ***p < 0.001.

**Suppl. Figure 3.**
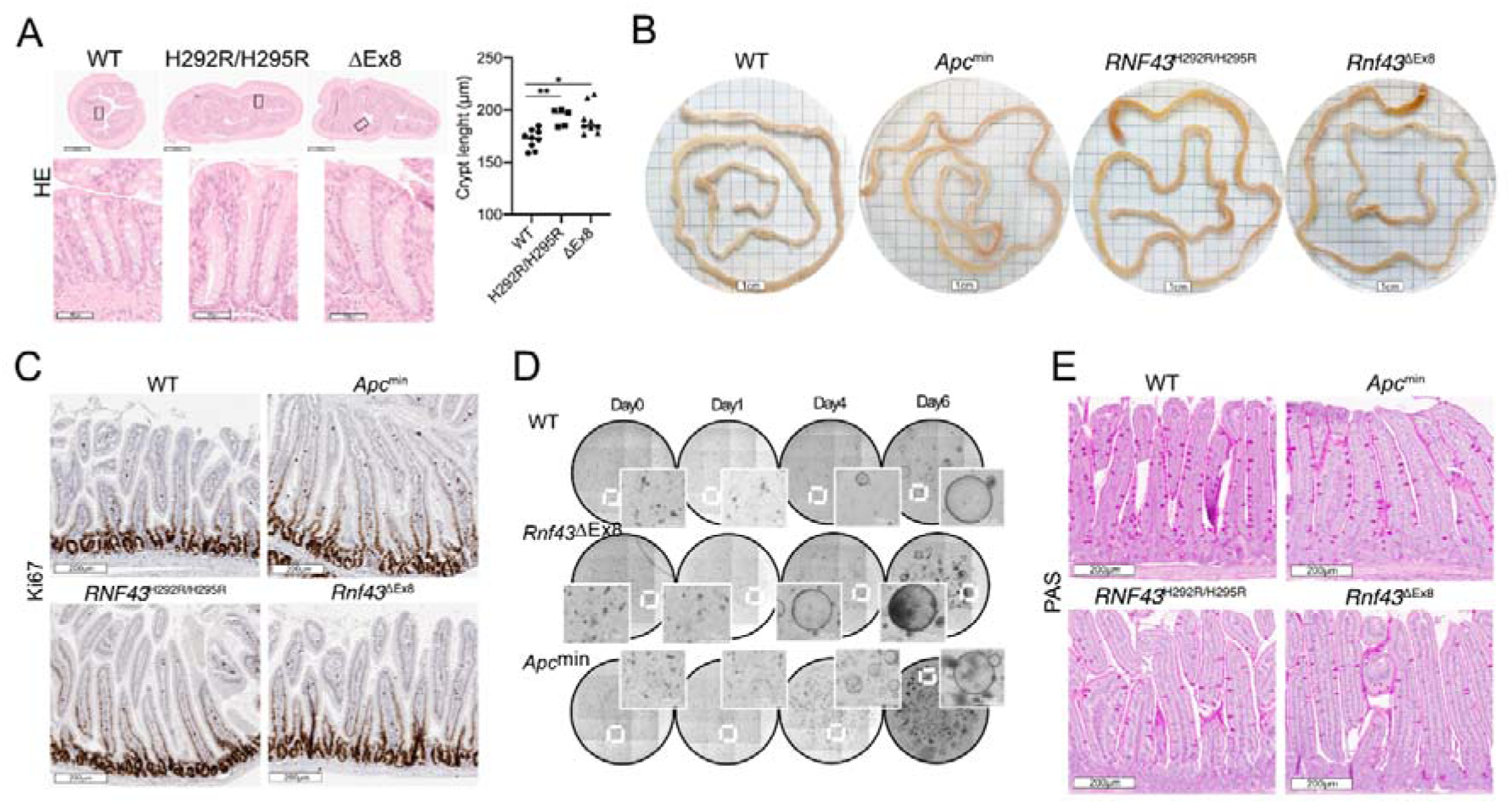
Mutations in *Rnf43* alter intestinal goblet cell homeostasis and microbiota composition. (A) Representative H&E-stained colorectal transections with zoomed-in crypt length (left), and crypt length quantification (right) in *Rnf43* mutant versus WT mice. (B-E) Analysis of young (3 months old) RNF43^H292R/H295R^, *Rnf43*^ΔEx8^ mice, *Apc*^min^ and WT mice in C57BL/6 background. (B) Representative intestinal appearance. (C) Representative Ki67-stained small intestine. (D) Density and size of colorectal organoids at the indicated days post manufacturing. (E) Representative PAS-stained small intestine. Each symbol represents one mouse. Bars indicate median. Ordinary one-way ANOVA with Tukey’s multiple comparisons correction was used for normal distribution, otherwise Kruskal-Wallis-Test with Dunn’s multiple comparisons correction, unless indicated differently. *p < 0.05, **p < 0.01.

**Suppl. Figure 4.**
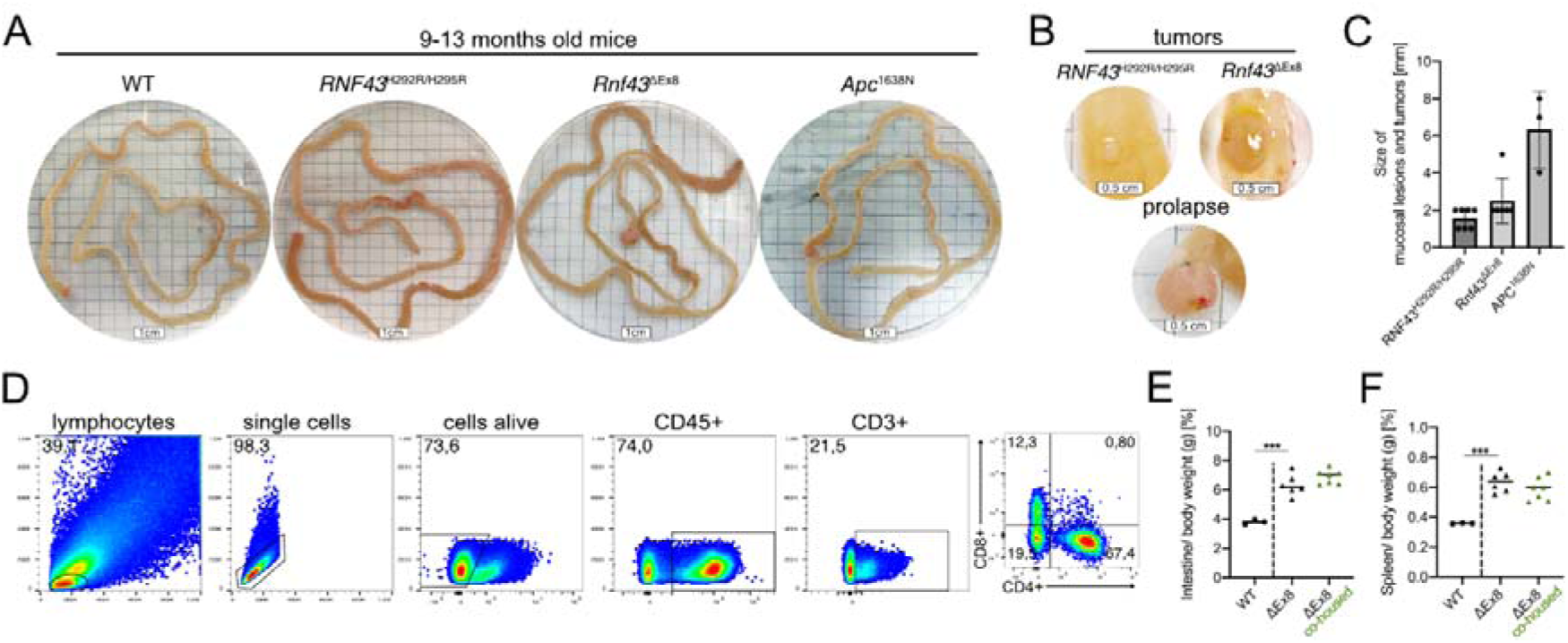
Mutations in *Rnf43* are sufficient to induce spontaneous intestinal inflammation with subsequent cancer development. (A-D) Intestinal phenotype of aging (9-13 months old) *Rnf43* mutant, *Apc*^1638N^ and WT mice in C57BL/6 background. Representative (A) intestinal reddening and (B) spontaneous intestinal tumor and rectal prolapse development in *Rnf43* mutant mice. (C) Tumor size in aging *Rnf43* mutant (*RNF43*^H292R/H295R^ n = 7, *Rnf43*^ΔEx8^ n = 6) versus *Apc*^1638N^ (n = 3) mice. (D) Flow cytometry gating strategy for lymphocytes. (E, F) Intestinal weight (E) and spleen weight (F) to body weight ratio in aging non-co-housed and co-housed *Rnf43*^ΔEx8^ mice. Each symbol represents one mouse. Bars indicate median. Ordinary one-way ANOVA with Tukey’s multiple comparisons correction was used for normal distribution, otherwise Kruskal-Wallis-Test with Dunn’s multiple comparisons correction, unless indicated differently. ***p < 0.001.

**Suppl. Figure 5.**
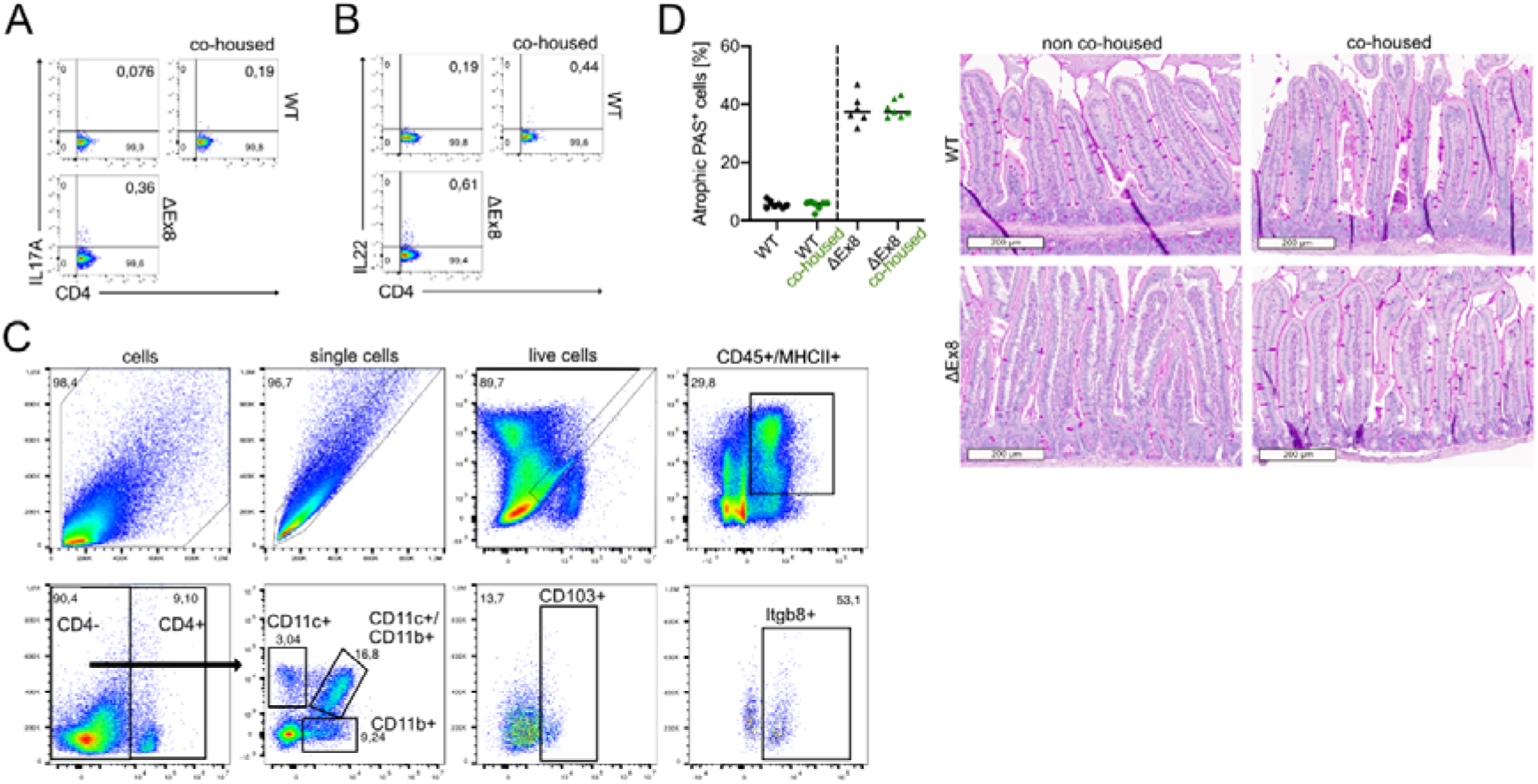
Epithelial inactivation of *Rnf43* causes reduced goblet cell density that is associated with DNA-damaging and Th17/Th22-inducing microbiota. (A, B) Representative flow plots of small intestinal CD4^+^ T cells of co-housed and non-co-housed WT mice releasing (A) IL-17A and (B) IL-22 after stimulation with PMA/Iono. (C) Flow cytometry gating strategy of ITGB8^+^ DCs. (D) Percentage and representative pictures of atrophic PAS^+^ goblet cells in the small intestine.

